# Characterizing the extracellular matrix transcriptome of cervical, endometrial, and uterine cancers

**DOI:** 10.1101/2022.04.04.486998

**Authors:** Carson J. Cook, Andrew E. Miller, Thomas H. Barker, Yanming Di, Kaitlin C. Fogg

**Author notes:** Address for correspondence: Kaitlin C. Fogg, School of Chemical, Biological, and Environmental Engineering Oregon State University, Corvallis, OR 97330.

## Abstract

Increasingly, the matrisome, a set of proteins that form the core of the extracellular matrix (ECM) or are closely associated with it, has been demonstrated to play a key role in tumor progression. However, in the context of gynecological cancers, the matrisome has not been well characterized. A holistic, yet targeted, exploration of the tumor microenvironment is critical for better understanding the progression of gynecological cancers, identifying key biomarkers for cancer progression, establishing the role of gene expression in patient survival, and for assisting in the development of new targeted therapies. In this work, we explored the matrisome gene expression profiles of cervical squamous cell carcinoma and endocervical adenocarcinoma (CESC), uterine corpus endometrial carcinoma (UCEC), and uterine carcinosarcoma (UCS) using publicly available RNA-seq data from The Cancer Genome Atlas (TCGA) and The Genotype-Tissue Expression (GTEx) portal. We hypothesized that the matrisomal expression patterns of CESC, UCEC, and UCS would be highly distinct with respect to genes which are differentially expressed and hold inferential significance with respect to tumor progression, patient survival, or both. Through a combination of statistical and machine learning analysis techniques, we identified sets of genes and gene networks which characterized each of the gynecological cancer cohorts. Our findings demonstrate that the matrisome is critical for characterizing gynecological cancers and transcriptomic mechanisms of cancer progression and outcome. Furthermore, while the goal of pan-cancer transcriptional analyses is often to highlight the shared attributes of these cancer types, we demonstrate that they are highly distinct diseases which require separate analysis, modeling, and treatment approaches. In future studies, matrisome genes and gene ontology terms that were identified as critical for predicting patient survival or cancer stage can be evaluated as potential drug targets and incorporated into *in vitro* models of disease.

## Introduction

Gynecological cancers are a major cause of cancer related death for women worldwide (Saso *et al*., 2011; Felix and Brinton, 2018; Torre *et al*., 2018; Arbyn *et al*., 2020). The most prevalent types include cervical, endometrial, and ovarian cancers. An estimated 91% of cervical cancer occurrences in the United States take place when the cells of the cervix are exposed to certain strains of human papilloma virus (HPV) (Castellsagué, 2008; Centers for Disease Control, 2021). Cervical cancer rates have been reduced in countries with robust healthcare systems due to improved screening and HPV vaccination; however, mortality and infection rates remain high – cervical cancer is the 2^nd^ leading cause of cancer-related death in women under 40 (Funston *et al*., 2018; Arbyn *et al*., 2020; Siegel *et al*., 2021). Endometrial cancer, the most common type of uterine cancer, forms in the lining of the uterus (Henley *et al*., 2018). Endometrial cancer is the most common gynecological cancer in the United States. Prevalence has increased by over 50% in recent decades, and survival rates for late-stage diagnosis are less than 15% (Saso *et al*., 2011; Funston *et al*., 2018). Ovarian cancer is the fifth most common cause of cancer related death in women in the US, with four out of five patients diagnosed with late-stage disease, leading to poor prognosis (Funston *et al*., 2018). While rare, uterine carcinosarcomas are a highly aggressive form of uterine cancer that form in the muscle lining of the uterus (Cherniack *et al*., 2017). Uterine carcinosarcomas account for under 5% of uterine tumors but are responsible for 30% of uterine cancer deaths – statistics which have not improved in several decades (Matsuo *et al*., 2016). Overall, while gynecological cancers are related by their involvement in the reproductive tract, each gynecological cancer originates in a unique tissue and has a unique pathology (Centers for Disease Control, 2020).

Increasingly, the matrisome, a set of proteins that form the core of the extracellular matrix (ECM) or are closely associated with it, has been demonstrated to play a key role in cancer progression, influencing epithelial-mesenchymal transition, angiogenesis, and metastasis (Hynes and Naba, 2012; Naba *et al*., 2012; Naba, Hoersch and Hynes, 2012; Mouw *et al*., 2014; Lim *et al*., 2019; Yadav *et al*., 2020). However, in the context of gynecological cancers, the gene expression of the matrisome has not been well characterized. A holistic, yet targeted, exploration of the tumor microenvironment is critical for better understanding the progression of gynecological cancers, identifying key biomarkers for cancer progression, establishing the role of gene expression in patient prognosis, and assisting in the development of new targeted therapies. It is also a key step in the construction of 3-dimenaional *in vitro* gynecological tumor models (Zhang *et al*., 2020), which could help accelerate drug screening and other research (Langhans, 2018; Wardwell-Swanson *et al*., 2020). In short, a deeper understanding of the transcriptome characteristics of the gynecological tumor matrisome has the potential to revolutionize the way gynecological cancers are understood and treated.

Publicly available datasets such as The Cancer Genome Atlas (TCGA) (Tomczak, Czerwińska and Wiznerowicz, 2015) and The Genotype-Tissue Expression (GTEx) database (Aguet *et al*., 2017) make it possible to do large-scale analyses of the transcriptomic profiles of different cancer types and tissues. Numerous studies have used these data to investigate overlapping expression patterns and enrichments between cancer types, find common features and subtypes, and explore novel techniques for analyzing these data (Cline *et al*., 2013; Khoury *et al*., 2013; Peng *et al*., 2015; Cestarelli *et al*., 2016; Dai *et al*., 2019; Clayton *et al*., 2020). Gynecological cancers have been included in pan-cancer studies (Berger *et al*., 2018; Lim *et al*., 2019; Parris, 2020), but often with the goal of highlighting shared attributes or analyzing them alongside breast cancer. With TCGA and other datasets, targeted analyses of gynecological cancers have been performed (Braun *et al*., 2013; Dellinger *et al*., 2016; Jha *et al*., 2017; M. Wang *et al*., 2018; Raffone *et al*., 2019; Zhang and Wang, 2019; Travaglino *et al*., 2020). These studies engage in analyses as specific as investigating the role of *L1CAM* in endometrial cancer progression (Dellinger *et al*., 2016), and as broad as analyzing expression, copy number variation, somatic mutation, and promoter methylation to identify pathways of interest across gynecological cancers (Jha *et al*., 2017). However, the need remains to conduct targeted analyses of the matrisome-level gene expression of individual gynecological cancers and explore their unique and shared attributes.

In this study, we used a combination of statistical methods and machine learning approaches to analyze bulk RNA sequencing (RNA-seq) and clinical patient data from a unified dataset of TCGA and GTEx (Q. Wang *et al*., 2018), allowing us to compare the matrisome gene expression of gynecological tumors to their corresponding healthy tissues. We explored the matrisome of four primary gynecological cancer types in the TCGA dataset: (1) Cervical squamous cell carcinoma and endocervical adenocarcinoma (CESC), (2) Ovarian serous cystadenocarcinoma (OV), (3) Uterine Corpus Endometrial Carcinoma (UCEC), and (4) Uterine Carcinosarcoma (UCS). Additional TCGA cancer cohorts, breast invasive carcinoma (BRCA), colon adenocarcinoma (COAD), testicular germ cell tumors (TGCT), and prostate adenocarcinoma (PRAD) were included as additional points of comparison. We then performed inferential analyses of the CESC, UCEC, and UCS cohorts to characterize the dysregulation of the tumors’ respective matrisome expression. Unfortunately, OV was excluded from these analyses due to the lack of sufficient readily available bulk RNA-Seq data for normal tissue samples to enable statistical inference. We identified differentially expressed genes (DEGs) and differentially expressed matrisome genes (DEMGs), individual matrisome genes and matrisome gene network modules which had inferential significance for patient survival and FIGO stage (the standard metric set by Fédération Internationale de Gynécologie et d’Obstétrique (FIGO) and used by pathologists when classifying gynecological cancer stage (Freeman *et al*., 2012)). We then used the DEGs, DEMGs, and the stage and survival significant matrisome genes to identify enriched gene ontology (GO) terms (groups of functionally related genes found to be overrepresented among gene sets of interest using enrichment analysis) (Ashburner *et al*., 2000). Cervical cancer, endometrial cancer, and uterine carcinosarcomas each had unique matrisome dysregulation profiles, highlighting how critical it is that gynecological cancers be viewed as highly distinct pathologies. Furthermore, our approach to identify matrisome genes and GO terms of interest can be applied to other types of cancer as well as other pathologies.

## Results

### Matrisome gene expression is selectively dysregulated in gynecological cancers is selectively dysregulated in gynecological cancers

To investigate the overall importance of the matrisome in characterizing tumor tissues, we performed differential gene expression (DGE) analysis on the full set of genes for each gynecological cancer cohort in our dataset *n_genes_*= 20,242), comparing tumor tissue to normal tissue (CESC: *n_normal_* = 13; *n_tumor_* = 259; UCEC: *n_normal_* = 105, *n_tumor_* = 14; UCS: *n_normal_* = 105, *n_tumor_* = 47). Functional enrichment analysis was then performed on the sets of DEGs to determine whether matrisome GO terms were enriched among the DEGs. We found large numbers of DEGs in all gynecological cancers: 7,652, 8,229, and 7,646 in CESC, UCEC, and UCS, respectively (**Figure 1A** and **Table 1**). We also found that a high proportion of matrisome genes were differentially expressed (**Figure 1B** and **Table 1**).

**Figure 1.**
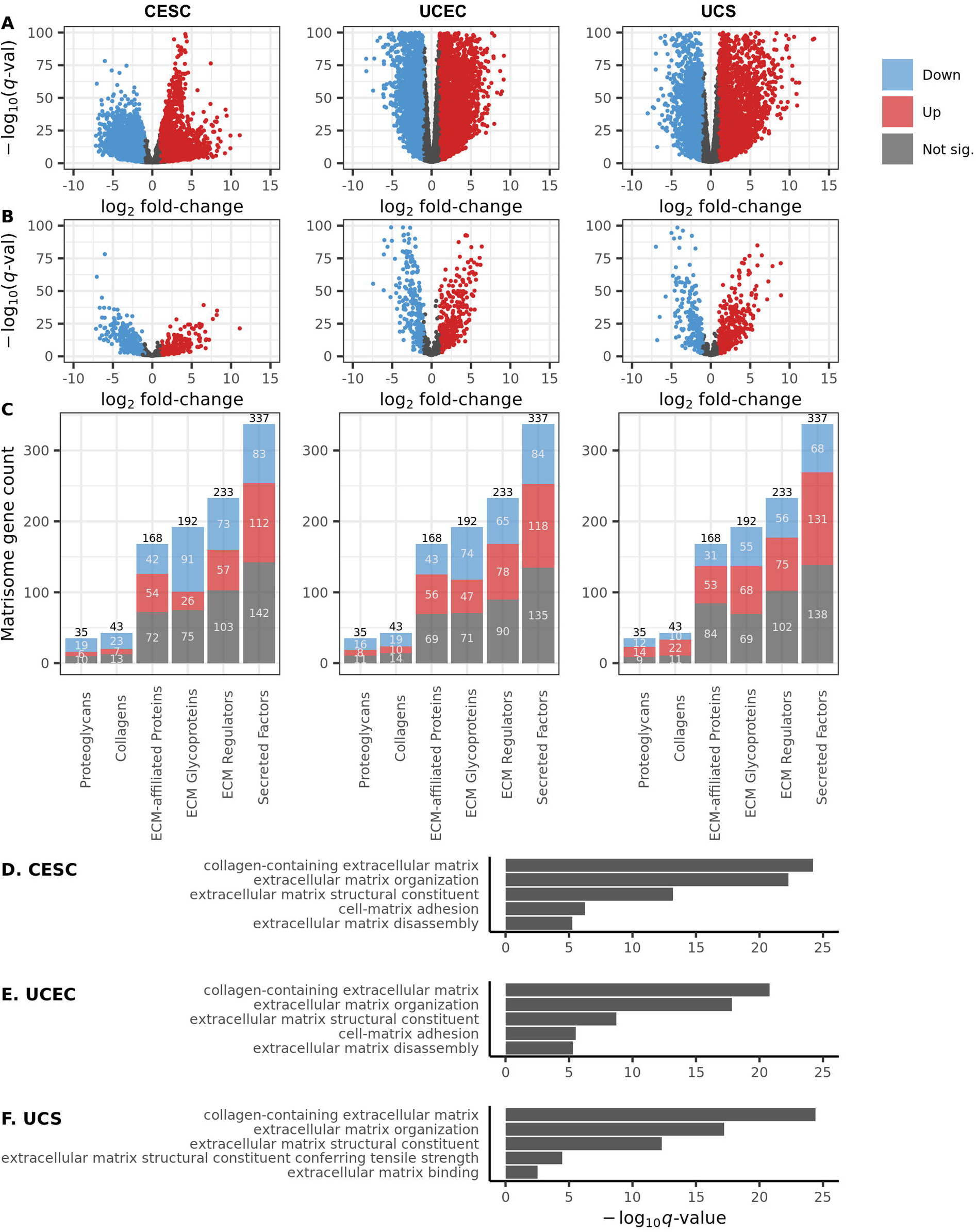
Differential gene expression and functional enrichment analysis. Volcano plots showing gene upregulation and downregulation for the cervical (CESC), endometrial (UCEC), and uterine (UCS) cohorts in **(A)** all genes and **(B)** matrisome genes only. **Observations** with – log_10_ *q*-values of ≥ 100 were excluded from visualization. **(C)** Breakdown of differential *expression* among matrisome genes by matrisome category. Gene expression color-coded by differential expression direction (blue = downregulated, red = upregulated, gray = no significant change). Sample sizes: CESC (*n_normal_* = 13, *n_tumor_* = 259), UCEC (*n_normal_* =105, *n_tumor_* = 141), and UCS (*n_normal_* = 105, *n_tumor_* = 47). Top 5 most enriched ECM-related gene ontology (GO) results for **(D)** CESC, **(E)** UCEC, and **(F)** UCS.

After DGE analysis had been performed, differentially expressed matrisome genes (DEMGs) were stratified using matrisome categories defined by Naba *et al*. (Naba *et al*., 2012) (**Figure 1C**, **Table S1**). The established matrisome categories are proteoglycans, collagens, ECM-affiliated proteins, ECM glycoproteins, ECM regulators, and secreted factors. While over half of the genes in all matrisome categories were differentially expressed in each cohort, collagens and proteoglycans were the most dysregulated in all three cancers (**Figure 1C**, **Table S1**). This is likely due, at least in part, to the fact that collagens and proteoglycans are constituents of the core matrisome and are more likely to be found in all tissues than the ECM-related matrisome genes (Naba *et al*., 2012). Additionally, we observed that ECM-related GO terms were consistently enriched among each cancer type (**Figure 1D-F**). In each cohort, the matrisome genes present in the unified dataset *n_mat.genes_* = 1,008) were approximately 20% more likely to be differentially expressed than the general gene population (**Table 1**). For example, in CESC, 38% of total genes were differentially expressed compared to 59% of matrisome genes. This strongly reinforces the idea that matrisome gene expression is critical for characterizing cancer pathology.

Next, we sought to demonstrate that the expression of the 1008 matrisome genes present in the unified dataset was sufficient for capturing a large amount of the information about the three gynecological cancers (CESC, UCEC, and UCS), even though the full matrisome constitutes only 4% of the human proteome (Naba *et al*., 2012). To this end, we trained elastic net penalized logistic regression (Zou and Hastie, 2005) models on the variance-stabilizing transformed (VST) unified matrisome expression data for each gynecological cancer. Because the numbers of tumor and normal observations were imbalanced in each cohort, especially in CESC (**Table S2**), balanced classification accuracy was used to assess model performance because of its ability to penalize poor performance on minority classes (Brodersen *et al*., 2010). The classifiers were able to attain perfect or near-perfect cross-validated balanced accuracy scores in all three cohorts when distinguishing between tumor and normal samples (**Table S2**). Given the performance of these models, the broad enrichment of ECM GO terms among DEGs in each cohort, and the widely acknowledged tissue-specific characteristics of ECM (Frantz, Stewart and Weaver, 2010; Hynes and Naba, 2012), it is clear that matrisome gene expression profiles capture valuable gynecological cancer-specific information.

### Gynecological tumor heterogeneity

To demonstrate tumor heterogeneity between gynecological cancer matrisome expression profiles, we used the VST harmonized TCGA matrisome expression data (NCI Genomic Data Commons, 2021) to perform clustering and dimensionality-reduced data visualizations. First, we computed centroids for each cancer type, constituted of gene-wise median matrisome gene expression values in each cancer, and identified 25 samples in each cancer which had minimum L^1^ (gene-wise absolute value) distance from these centroids. Correlative hierarchical clustering on these representative samples showed clear matrisome heterogeneity between gynecological tumors, though there were a few representative UCS and UCEC samples which correlated with the OV samples more closely than the rest of their cohort (**Figure 2A**). Matrisome gene expression of the gynecological cancer cohorts was then compared to selected non-gynecological cancer TCGA sample cohorts: BRCA (breast invasive carcinoma), COAD (colon adenocarcinoma), TGCT (testicular germ cell tumors), and PRAD (prostate adenocarcinoma), all of which share certain characteristics of gynecological cancers. Correlative hierarchical clustering was performed again, and the results showed instances in which the matrisome gene expression profiles of gynecological tumors correlated more closely with non-gynecological tumors than with other gynecological tumors (**Figure 2B**). For example, many of the representative UCS samples clustered more closely with TGCT than other the gynecological cancer types, and CESC showed tighter clustering with COAD than the other gynecological types. This indicates that, at the level of matrisome gene expression, gynecological cancers do not exhibit a unified phenotype.

**Figure 2.**
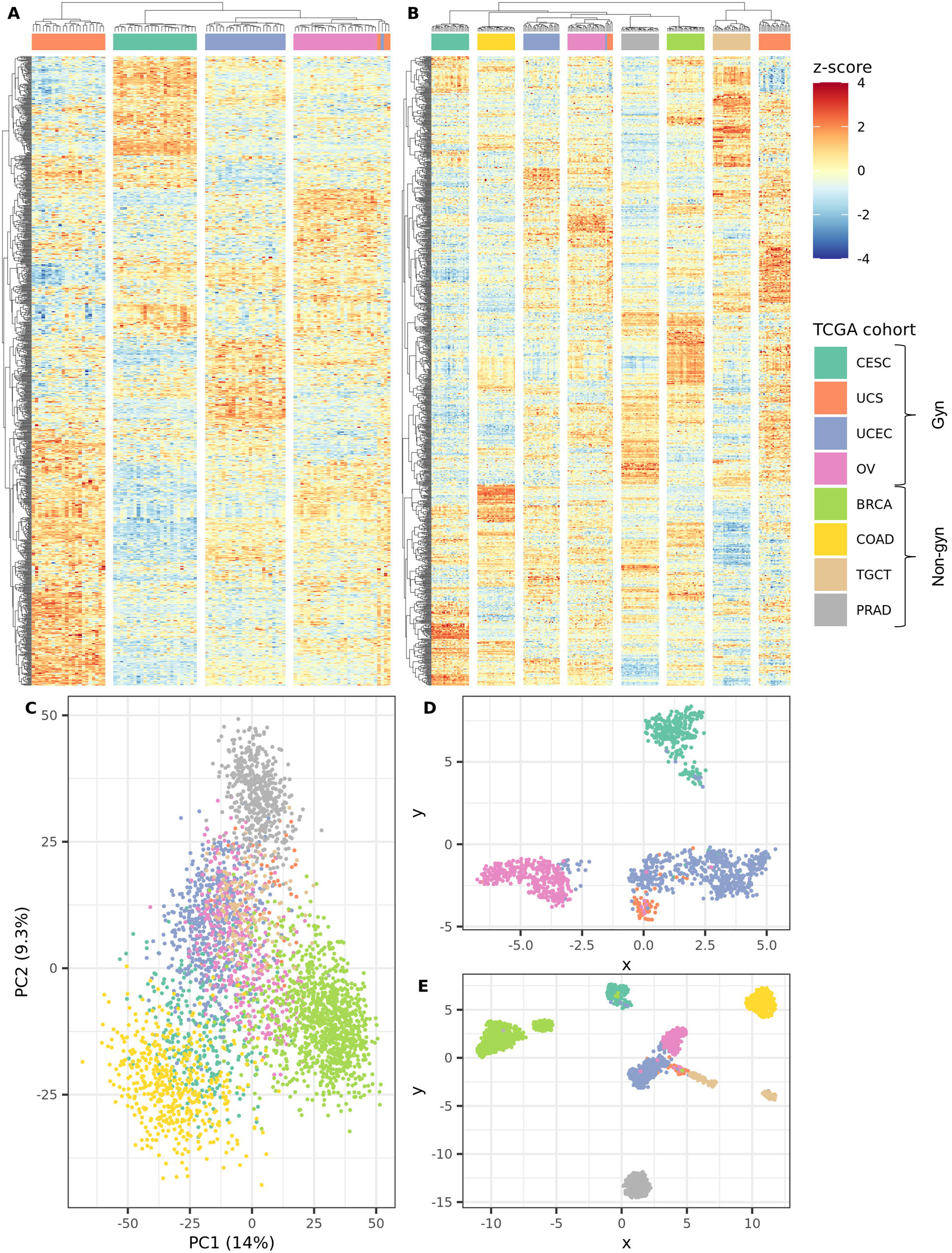
Matrisome heterogeneity across gynecological cancers. Correlative hierarchical clustering heatmaps of matrisome genes (rows) and samples (columns) using 25 representative samples from **(A)** gynecological (gyn) cancers only and **(B)** gynecological and non-gynecological (non-gyn) cancers together. **(C)** Scatterplot of principal components 1 and 2 (14% and 9.3% variance explained, respectively) for gynecological and non-gynecological cancers. UMAP dimensionality reduction performed on **(E)** gynecological cancers only and on **(F)** gynecological and non-gynecological cancers together.

These differences were further explored through principal component analysis (PCA) and Uniform Manifold Approximation and Projection (UMAP). PCA largely failed to clearly separate subpopulations (**Figure 2C**, **Figure S1**), likely because of its inability to perform non-linear transformations. To remedy this, we applied UMAP (McInnes, Healy and Melville, 2020), which can capture non-linear information in its approach to dimensionality reduction. UMAP further demonstrated that matrisome gene expression was highly distinct between gynecological cancer types. While there was evidence of small subgroup overlaps, overall, the different gynecological cancer cohorts formed quite distinct clusters (**Figure 2D**). Though the distances between subpopulations identified by UMAP are not directly interpretable due to the nature of the algorithm, CESC samples clustered rather tightly and were mostly separated from the other gynecological cancers while UCEC and UCS had a large amount of overlap. Like CESC samples, OV samples clustered mostly among themselves, but overlaps were observed with UCEC samples (**Figure 2D**). Again, the matrisome gene expression of the gynecological cancers were then compared to non-gynecological cancers BRCA, COAD, TGCT, and PRAD (**Figure 2E**). Interestingly, we observed that there was significant overlap and contiguity between UCS and TGCT. Additionally, similar to the UMAP analysis of gynecological cancers alone, a small subgroup of UCEC samples overlapped with OV samples and several UCS samples were interspersed among the UCEC cohort (**Figure 2D, E**). Importantly, breast cancer was as visually distinct from gynecological cancers as prostate and colon cancers (**Figure 2E**, **Figure S1**). The fact that breast cancer samples cluster so tightly away from gynecological cancer samples while testicular cancer samples have surprising contiguity with uterine carcinosarcoma samples implies that matrisome gene expression reveals key similarities and differences between how tissues experience cancer invasion which might be missed using traditional sex-based health approaches to cancer analysis. Also, given that ECM is highly tissue-specific, the overlapping subgroups may represent instances in which the respective cancers have remodeled or dedifferentiated ECM tissues in a similar way. Taken together, these data demonstrate that there are instances where the matrisome gene expression patterns of gynecological cancers may be more similar to non-gynecological cancers than each other, and that, while overlapping subpopulations are observable, the gynecological cancers form distinct clusters.

### Univariable and multivariable analysis to assess predictive power for cancer stage and patient survival

We developed a multi-faceted approach to investigate the relationship of individual matrisome gene signatures to cancer stage and patient survival in each gynecological cancer, using FIGO stage and patient survival as stratification variables. Analyses were performed using the unified gene expression data (Q. Wang *et al*., 2018) paired with TCGA clinical data. For analyzing FIGO stage-gene relationships, three approaches were used in each cohort: 1) gene-wise point-biserial correlation between genes and each FIGO stage (Kornbrot, 2005; Langfelder and Horvath, 2008), 2) pairwise differential gene expression analysis between samples from the different FIGO stages, and 3) multivariable L^1^ penalized multinomial regression (Schölkopf, Platt and Hofmann, 2007; Friedman, Hastie and Tibshirani, 2010) to create parsimonious models (inferentially significant models with minimal parameters) which yielded subsets of matrisome genes with explanatory power for tumor progression. A similar approach was used for exploring survival-gene relationships. Here the three techniques included: 1) gene-wise censored time screening (Cox, 1972; Langfelder and Horvath, 2008), 2) a gene-wise union of Kaplan-Meier and Cox proportional hazards (Cox PH) analysis (Kaplan and Meier, 1958; Cox, 1972), and 3) multivariable L^1^ penalized Cox PH (Goeman, 2010; Simon *et al*., 2011), which, like the FIGO L^1^ penalized multinomial regressors, created parsimonious models which yielded explanatory subsets of matrisome genes but this time with respect to patient survival.

Within the FIGO stage and survival analyses, each modeling approach asked a subtly different question, so the union of their respective results represents a more holistic set of matrisome genes which are important with respect to FIGO stage or patient survival. The pairwise FIGO stage differential expression analysis utilized all gene counts for cancer patients, and computed results were filtered down to only those genes contained within the matrisome. All other analyses utilized the VST cancer patient expression data for the matrisome genes. Matrisome genes which were found to have significance in terms of descriptive or inferential value via these methods were filtered based on DEMG status and then deemed model significant. We used the DEMGs as a filtration mechanism since we were primarily interested in those matrisome genes which are dysregulated in cancer overall *and* hold significance in some other respect (e.g., stage or survival significance). We did not pre-filter by DEMG status before our analyses to avoid biasing the results of those models which used global information (e.g., differential expression or L^1^ penalized models), or artificially decrease *p*- and *q*-values by reducing the number of tests. DEMGs which held univariable and/or multivariable significance with respect to FIGO stage were termed *stage model significant* while those which held univariable and/or multivariable significance with respect to patient survival were termed *survival model significant* (**Table S3**). Model significant DEMGs are matrisome genes which hold the most information about cancer stage and/or patient survival, and which may also serve as proxies for the impacts of co-expressed matrisome genes not captured through these analyses.

A stark difference was observed between CESC and the other two gynecological cancer cohorts with respect to multivariable FIGO stage analyses. L^1^ multinomial regression yielded a large set of stage model significant DEMGs for CESC *n* = 105), while it yielded much smaller sets in, UCEC and UCS (*n* = 13 and *n* = 38, respectively) (**Table S3**). The number of univariable significant DEMGs was more similar between the cohorts. This could point to a more diffuse relationship between matrisome genes and cancer stage in CESC, with many different groups of genes impacting cancer stage in different ways, rather than a few key drivers/proxy genes. In our survival analysis, all gynecological cancers yielded a similar number of multivariable results. For univariable survival analysis, however, CESC had far more significant DEMGs than UCEC or UCS – for example, univariable Kaplan-Meier and Cox PH analysis yielded 19 significant DEMGs in CESC, but only 1 and 2 in UCEC and UCS, respectively (**Table S3**).

For stage model significant DEMGs, no genes were shared by all cancers, but we did identify 14 genes shared between CESC and UCEC, 12 genes between CESC and UCS, and 7 genes between UCEC and UCS. Though no stage model significant DEMGs were shared between all three cancers, we did note the presence of SERPIN-family members in results for each of the three cancer cohorts. Further, CESC and UCEC share a strong presence of immunomodulatory cytokines and chemokines which have previously been tied to cancer progression in several cancer types (Mathur *et al*., 2003; Massagué, 2008; Bruchim, Sarfstein and Werner, 2014). However, the results for the three cohorts were more distinct than similar. CESC demonstrated the most depth (quantity of genes within the same family) of dysregulation across the gene families which were found among stage model significant DEMGs. CESC stage model significant DMEGs were enriched for genes related to hedgehog signaling (*HHIP, IHH, and SHH*), and the mucin family of *O*-linked glycoproteins (*MUC2/3A/5AC/6/12*). Several ECM protease families were also highly enriched among these genes – the *ADAM* family (*ADAM11*/*20*/*22*/*29*/*8*), *MMP*s (*MMP7/8/9/10/13/17/25/27*), and *ELANE*, which codes for neutrophil elastase. In contrast, UCEC demonstrated little depth or breadth of family representation, though stage model significant DEMGs were observed to contain multiple ECM binding regulators (*HRG*, *OTOG*, and *PCOLCE2*) and extracellular enzymes (*HYAL3* and *PLG*). We also noted that UCEC exhibited a unique overrepresentation of members of the TGF-β super family, specifically towards the growth differentiation factor subfamily. UCS showed a large breadth of family representation, but little depth within families. Stage model significant DEMGs were found within the cystatins (*CST6* and *CSTB*), *S100* family of Ca^2+^-binding proteins (*S100A7/8/9/14* and *S100B/P*), and the glycosaminoglycan-binding proteins *NCAN*, *NELL2*, and *ECM2*. Together, these data suggest that no singular DEMGs or set of DEMGs are constitutively tied to FIGO stage, but rather tissue-specific mechanisms employed by each cancer subtype may modulate oncogenesis and increasing FIGO stage.

For survival significant DEMGs, we identified a single gene with overlap between CESC and UCEC (*RSPO2*, a member of the WNT signaling pathway). No other genes were shared among the three cancers. CESC again demonstrated the most depth and breadth of represented gene families among DEMGs in this category. CESC was characterized by survival model significant DEMGs largely related to immune recruitment and immunomodulatory signaling (*CCL25*, *CSF2*, CXCL2/3, *IL-1b*, *LIF*, *TNF*, *TNFSF15*, and *TGFB1*) as well as regulators of matrix protein processing (*MMP1*/*3*, *TLL1*, and *PLOD2*). UCEC survival model significant DEMGs were characterized by less depth and breadth than the stage model significant DEMGs and lacked clear overrepresentation of a specific gene family. Survival model significant DEMG sets for UCEC included ECM components (*CSPG4*, *COL11A1*, *FREM3*) alongside representation for genes ascribed to regulation WNT signaling (*RSPO2 and WIF1*) and immune cell adhesion (*CLEC12B*, *MEGF10*, *MUC2*). For survival, UCS had the least depth and breadth, but demonstrated dysregulation and prognostic value within mucins (*MUC12*/*17*) and ECM regulation/organization (*TIMP4*, *COL5A3*, *HAPLN2*, and *NCAN*). These data support a framework in which genes predictive of survival are cancer-specific and not preserved amongst the various gynecological tissues of origin.

### Weighted gene correlation network analysis

To examine clusters of co-expressed genes that were significantly related to tumor stage and patient survival, we used weighted gene correlation network analysis (WGCNA) (**Figure 3**). WGCNA uses unsupervised clustering on topological overlap measures (TOMs) between genes to construct gene modules which correlate with sample traits of interest, which for our study were either FIGO stage or patient survival (Langfelder and Horvath, 2008). TOM is a measurement which is often used to approximate gene co-expression (Yip and Horvath, 2007), and module-trait relationships are typically assessed by correlating module eigengenes – the first principal component of a gene module – with the trait of interest (Langfelder and Horvath, 2008).

**Figure 3.**
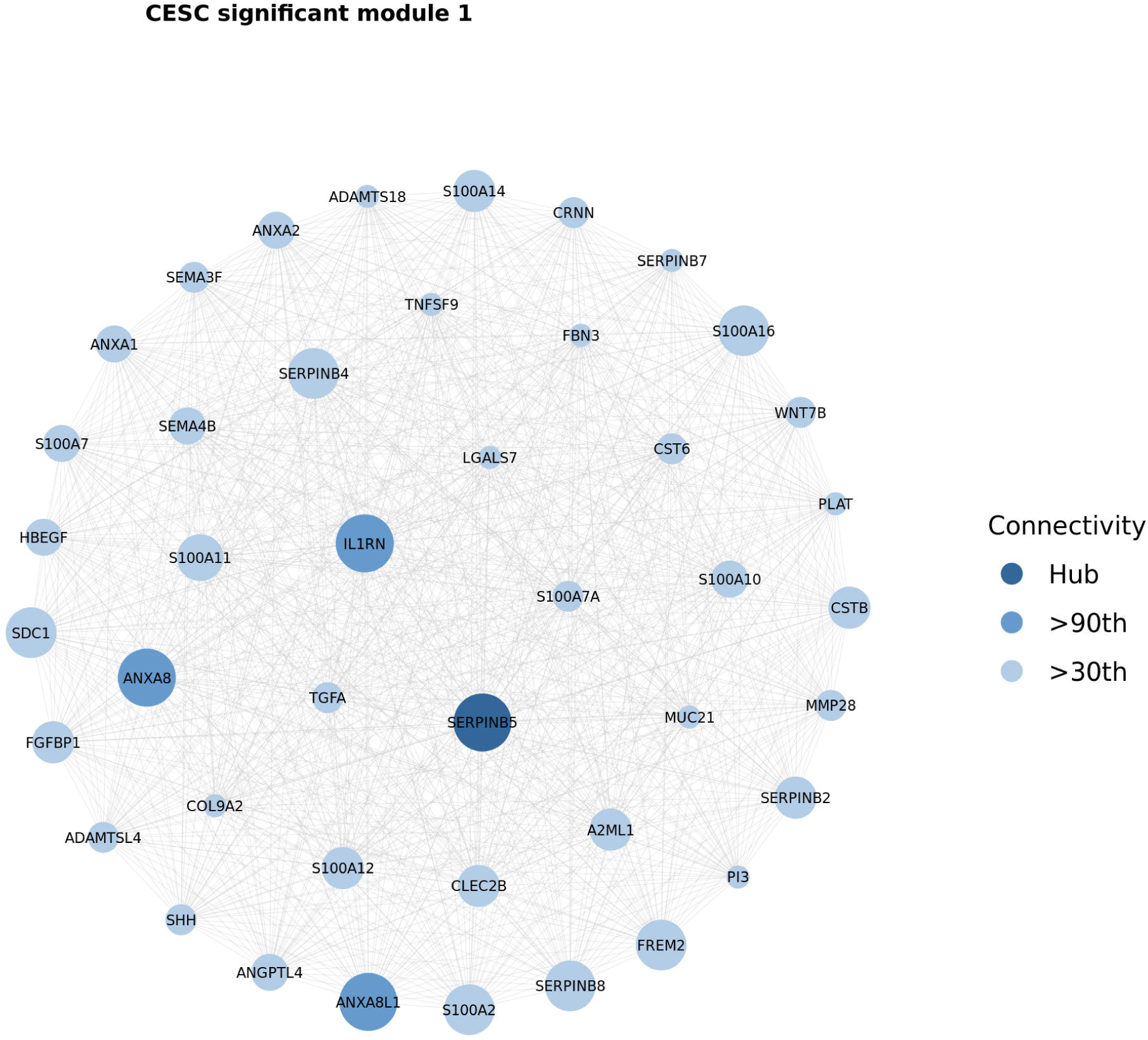
Weighted gene correlation network analysis. Representative visualization of a weighted gene correlation network analysis (WGCNA) module. This module represents DEMGs that were significantly correlated with FIGO stage in CESC. These genes were filtered, scaled, and shaded based on their connectivity as compared to the connectivity of their module’s hub gene, the most connected DEMG in the module. For visualization purposes, module DEMGs which were below the 30^th^ percentile of connectivity are not pictured, though they were utilized in our analyses. Module DEMGs which were in the 90^th^ percentile of connectivity to the other module genes are shaded darker than those below the 90^th^ percentile. Hub genes are shaded darkest. Connectivity was determined by the row-wise (gene-wise) sum of the module’s adjacency matrix.

WGCNA was performed on the VST unified matrisome expression data within each gynecological cancer cohort and separate tests were conducted identify modules and their constituent matrisome genes which were significantly correlated with FIGO stage or survival (Student asymptotic *q* < 0.05). We also explored the overall TOM profiles of the matrisome in all three gynecological cancers to compare the gene-gene interactions of the cancers. CESC matrisome gene expression demonstrated mean matrisome gene-wise TOMs far exceeding those observed in UCEC or UCS, implying that matrisome gene co-expression is more broadly distributed among matrisome genes in CESC compared to UCEC or UCS (**Figure S2** and **Table S4**). This, along with the results from the univariable and multivariable analyses, implied that CESC is characterized more by large matrisome gene ensemble behavior than expression levels of a few genes serving as proxies for other co-expressed genes.

For each cancer, the matrisome genes within FIGO stage significant network modules were filtered by the significance of their correlation with their respective module eigengenes (Student asymptotic *p* < 0.05), and then filtered based on whether they had been identified as DEMGs. These DEMGs were deemed *stage network significant*. In the CESC, UCEC, and UCS cohorts, 140, 159, and 129 DEMGs were shown to be stage network significant, respectively (**Table S5**). WGCNA did not identify any network modules with eigengenes significantly related to patient survival in any of the gynecological cancers examined (log-rank test *q* < 0.05).

WGCNA revealed differences between cohorts in terms of the most connected DEMGs within their respective WGCNA modules. In CESC, the most connected DEMGs in the three significant modules were: 1) serine (*SERPIN*s) and cysteine (cystatins; *CSTs)* protease inhibitors, S100- family immune modulators, and annexins; 2) mucins; 3) immunomodulatory factors (*S100* family, *LIF*, *CXCL*s, *CCL*s, *TNFSF13, and IL17C*) and additional ECM protease inhibitors (*CST3, ITIH4, SERPIN*s, *SLPI, and TIMP1*) (**Figure 3, S3A**). In UCEC, the most connected DEMGs in the four significant modules were: 1) proteases (*ADAMTS* family and *HABP2*) and protease inhibitors (*SERPINs*, *ITIH2,* and *PZP*); 2) hormones regulating growth and metabolism (*CSH1/2, INSL3,* and *GH2*) as well as numerous factors associated with fibrin clot formation and hemostasis; 3) TGF-β superfamily members and mediators of SMAD signaling; and 4) both structural collagens and mediators of ECM assembly/turnover (**Figure S3B**). UCS had three significant modules where the most connected DEMGs were: 1) Members and mediators of WNT, FGF, and TGF-β family signaling; and 2) ADAM and cathepsin protease families in addition to Ca^2+^-binding immunoregulatory factors from the S100 and annexin families. The third module in UCS did not provide any novel insights but did further reinforce trends seen in other modules including WNT, FGF, and TGF-β growth factor signaling, protease inhibitors, and hormone signaling (**Figure S3C**).

### Significant matrisome gene overlaps

The primary results produced by our analyses were sets of DEMGs which were either FIGO stage or survival significant (**Figure 4A**, **Table S6**). All cohorts had high levels of overlap between DEMGs and both stage significant and survival significant matrisome genes. Additionally, for all cohorts, the univariable and multivariable analyses yielded very different gene results than our network analyses. Only 42, 22, and 14 matrisome genes were found to be significant among both model and network significant DEMGs in CESC, UCEC, and UCS, respectively. While unsurprising, this demonstrates the value of combining findings from several univariable and multivariable analysis approaches with results network analyses, as they have complementary strengths and weaknesses. We also observed that focusing on the overlap of matrisome genes that were differentially expressed in tumor versus normal and either model or network significant genes dramatically narrowed the number of matrisome genes of interest (**Figure 4B**). With respect to tumor stage significance, we were primarily interested in matrisome genes which were dysregulated in gynecological tumors overall *and* were implicated in tumor progression, and this method enabled that analysis.

**Figure 4.**
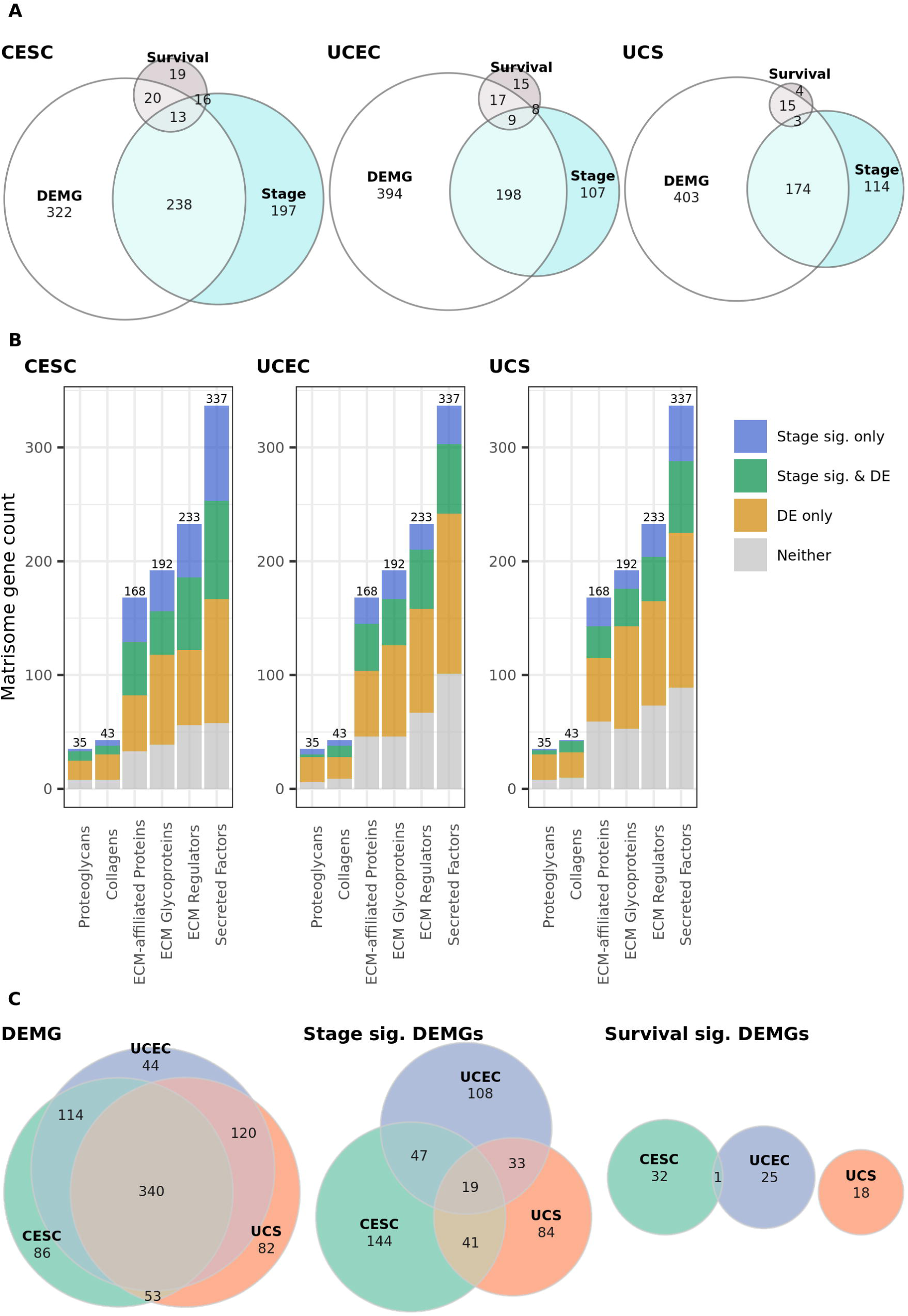
Significant matrisome gene overlaps. **(A)** Visualization of the overlaps between differentially expressed matrisome genes (DEMGs), stage significant matrisome genes (Stage), and survival significant matrisome genes (Survival) for each gynecological cancer cohorts. Stage or survival significant refers to all of the genes that were found by univariable, multivariable, or network analysis to be significant with respect to tumor stage. **(B)** Visualization of the overlap between differentially expressed (DE) and stage significant matrisome genes (Stage sig.) broken down by matrisome category for each gynecological cancer cohort. *Stage sig. only* refers to matrisome genes found by univariable or multivariable analysis or by network analysis to be significant with respect to tumor stage, but which were not identified as differentially expressed between tumor and normal tissue. *DE only* refers to matrisome genes differentially expressed between tumor and normal tissue but not identified as significant through univariable or network analysis. **(C)** Visualization of the overlaps between gynecological cohorts in terms of differentially expressed matrisome genes (DEMGs), stage significant DEMGs, and survival significant DMEGs.

Heterogeneity between the gynecological cancer cohorts in terms of stage or survival significant DEMGs was then visualized (**Figure 4C**). A striking amount of inter-cancer overlap was observed with respect to DEMGs, with 340 of 839 unique DEMGs shared between all gynecological cancer cohorts. However, the amount of overlap with respect to stage significant DEMGs was much smaller, with only 19 out of 476 unique matrisome genes shared by all cohorts. Survival significant DEMGs had limited overlap among the 76 unique survival significant DEMGs, with UCS having no overlap with the other two cohorts, and CESC and UCEC sharing only one survival significant DEMG, *RSPO2*, a member of the R-spondin protein family which codes for a ligand involved in Wnt signaling. Interestingly *RSPO3*, another member of the R-spondin protein family which codes for a protein linked to Wnt signaling regulation, was also shared by CESC and UCEC among stage significant DEMGs. These overlap summaries imply that the underlying mechanisms for tumor progression in the three cohorts are distinct yet have significant overlap, while the underlying mechanisms impacting patient survival are highly distinct.

DEMGs that were stage significant in all three gynecological cancer types included *PRL* (prolactin, critical to mammary gland development and lactation through regulation of estrogen receptors (Lopez Vicchi and Becu-Villalobos, 2017); upregulated in CESC and UCEC, but downregulated in UCS), *CCL28* (promotes angiogenesis in response to hypoxia (Facciabene *et al*., 2011); upregulated in all cancers), *ADAMTSL4* (positively regulates fibrillin-1 which regulates latent *TGFb* sequestration (Apte, 2009); downregulated in all three cohorts), and *MUC13* (shares *EGFR* domain with *MUC1*, which promotes *MUC1*-*bCat* interaction (Putten and Strijbis, 2017); upregulated in all three cohorts). Families of genes that were among the stage significant DEMGs in each cancer type included those related to immune signaling, ECM modification/regulation, growth factor signaling, and cell adhesion regulation. Some pair-wise overlaps of interest among stage significant DEMGs include the following: 1) transcripts encoding laminins and growth differentiation factor family members (*TGF-*β superfamily proteins) were significant in both UCS and UCEC, 2) MMPs, FGFs, and cytokines/chemokines(*IL/CCL/CXCL* transcripts) were found to be significant in CESC and UCS, and 3) SERPIN, C3 & PZP, and ITIH protease inhibitor families as well as c-type (CLEC) and galactoside-binding (LGALS) lectins were found to be significant in CESC and UCEC.

### Functional enrichment analysis

Lastly, we performed functional enrichment analysis, resulting in cellular component, biological process, and molecular function gene ontology (GO) classifications that were enriched among the genes found by our analyses (Gene Ontology Consortium, 2004). We identified GO terms which were enriched among stage or survival significant DEMGs and sorted them by tl-value (**Figure 5**). For stage significant DEMGs (**Figure 5A**), out of the top 10 most significantly enriched GO terms in each cohort (14 unique GO terms total), 7 were shared by all cohorts (extracellular matrix organization, extracellular structure organization, collagen-containing extracellular matrix, receptor ligand activity, signaling receptor activator activity, cytokine activity, and growth factor activity), a 50% full overlap. Examining all 974 GO terms that were significantly enriched in one or more cohort across all stage significant DEMGs, 221 were significantly enriched in all cohorts, a 23% overlap. The observed overlap indicates that despite the specific genes differing, the three cancers approach several of the same endpoints through different means.

**Figure 5.**
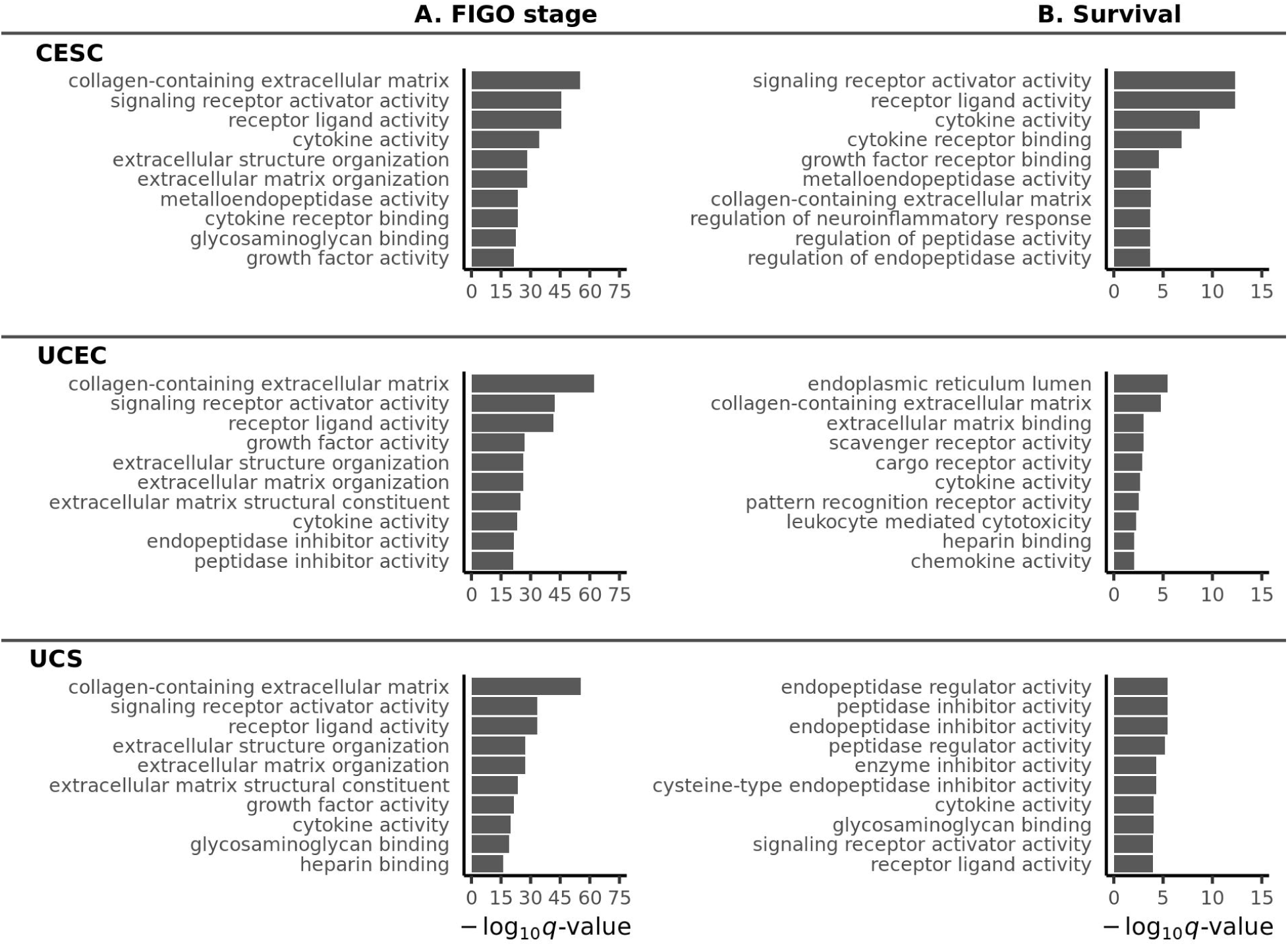
Functional enrichment of FIGO stage and survival significant DEMGs for all gynecological cancer types. Top 10 most significantly enriched gene ontology terms (sorted by tl-value) in each cancer cohort with respect to **(A)** FIGO stage significant differentially expressed matrisome genes (DEMGs) and **(B)** survival significant DEMGs. Value of – log_10_ (*q*) > 1.3 indicates significance (q < 0.05).

For survival significant DEMGs (**Figure 5B**), out of the top 10 most significantly enriched GO terms in each cohort (25 unique GO terms total), only 1 (cytokine activity) was shared by all cohorts, a 4% full overlap. Examining all 457 GO terms that were significantly enriched in one or more cohort across all survival significant DEMGs, only 8 were shared by all cohorts, a <2% full overlap. Taken together, these data indicate that the GO terms enriched among DEMGs related to patient survival are broadly distinct between CESC, UCEC, and UCS.

The GO classifications of DEMGS identified as stage significant using either univariable or multivariable analysis were then compared to those identified using network analysis. Among stage model significant DEMGs identified through univariable or multivariable analysis in CESC, the most highly enriched GO terms were related to ECM structure, ECM organization, ECM deposition and remodeling, cytokine activity, receptor binding, growth factor activity, glycosaminoglycan binding, and receptor ligand activity. Examining matrisome genes within FIGO stage significant network modules, CESC module 1 showed enrichment of collagen ECM and negative regulation of peptidase/endopeptidase/serine-type endopeptidase activity, along with enrichment of calcium-dependent protein binding and enzyme inhibition. In module 2, innate immune response, O-glycan processing and protein O-linked glycosylation, and stimulation of the C-type lectin receptor signaling pathway were significantly enriched. The most enriched GO terms in CESC module 3 were related to chemokine and cytokine activity, as well as neutrophil and granulocyte movement. Overall, CESC was characterized by widespread dysregulation in the ADAM/TS/TSL and MMP protein families which contribute to the enrichment of the metalloendopeptidase GO term.

Among stage model significant DEMGs in UCEC, the most highly enriched GO terms were related to ECM structure, RAGE receptor binding, negative regulation of ECM remodeling, glycosaminoglycan binding, and calcium-dependent protein binding. Examining matrisome genes within FIGO stage significant network modules, the four significant network modules in UCEC had some overlap but were also quite distinct in terms of enriched GO terms. In UCEC module 1, collagens with collagen ECM and peptidase/endopeptidase/serine-type endopeptidase inhibition were highly enriched. Analysis of module 2 identified enzyme inhibition, blood microparticles, collagen ECM, glycosaminoglycan binding, peptidase/endopeptidase/serine-type endopeptidase inhibition, and glycosaminoglycan binding as highly enriched. Among the genes in module 3, tumor necrosis factor receptor binding, cell adhesion, cytokine activity, and growth factor activity were all significantly enriched, as were BMP-related GO terms. In UCEC module 4, the most enriched GO terms were related to ECM organization/collagen and metallopeptidase/metalloendopeptidase activity. Overall, UCEC dysregulation, in contrast to CESC, was characterized by growth factor signaling – components of the *FGF* (*FGF10*/*12*), *TGF-*β superfamily (*TGF-*β/*GDF*/*BMP* signaling; *TGFb1*, *LTBP3*, *GDF6*, *GDF7*, *BMP5*, *BMPER*), *IGF* (*IGF1*, *IGFBP2*/*6*), and *WNT* (*WNT10A*, *WNT11*, *RSPO2*) were all well represented.

Among stage model significant DEMGs identified through univariate or multivariate analysis in UCS, the most highly enriched GO terms were related to ECM structure, ECM organization, ECM deposition and remodeling, enzyme inhibition, growth factor activity, cytokine activity, and negative regulation of proteolysis were the most enriched. Examining matrisome genes within FIGO stage significant network modules, module 1 showed enrichment of ECM structure/organization, semaphorin/plexin signaling, keratinocyte migration, and regulation of transmembrane receptor proteins. Module 2 showed significant enrichment of ECM collagen/structure/organization, cytokine activity, and growth factor activity. UCS module 3 showed significant enrichment of S100 protein binding, semaphorin receptor binding, glycosaminoglycan binding, and negative regulation of chemotaxis. Overall, UCS was characterized by matrix modifying enzymes (*LOXL3*, *MMP1*/*9*/*21*/*23B*, *CELA2A*/*B*, *CELA3A*/*B*), cell-surface proteoglycans (mucins *MUC2*/*13*/*15* and glypicans *GPC1*/*2*/*3*/*6*), and ECM structural components (collagens *COL19*/*20*/*22*/*23*/*25*/*28*/*A1* and laminins (*LAMA2*/*3*, *LAMB3*, *LAMC2*).

As there were no survival network significant modules identified using WCGNA, the survival model significant DEMGS identified through univariate or multivariate analysis were explored in relation to only the other types of cancers. Overall, survival significant DEMGs were characterized by a less diverse set of enrichments. In CESC, immunomodulatory cues were dominant, with MMPs and collagen processing proteins such as *PLOD2* and *ITIH1* also showing noteworthy levels of representation. In UCEC, enriched GO terms were characterized by heparin binding proteins and mediators of WNT pathway activation and signaling. Lastly, UCS enrichments were predominantly composed of glycosaminoglycan binding proteins with a bias towards hyaluronic acid and heparin binding factors.

## Discussion

In summary, we demonstrated that the matrisome is more dysregulated in gynecological cancer than genes at large and is highly critical for characterizing gynecological cancers. We reinforced the fact that gynecological cancers are highly heterogeneous and distinct from breast cancer at the matrisome level. Through a combination of statistical and machine learning methods, we identified differentially expressed matrisome genes and used univariable and multivariable modeling as well as network analysis to subcategorize these into those which were descriptively or inferentially significant with respect to either tumor stage or patient survival. We then determined GO terms which were significantly enriched within each of those sets of genes.

Finally, the results of our individual gene and gene network analyses as well as our enrichment analyses were used for two tasks. First, within each gynecological cancer cohort, we contrasted the genes and GO terms found to be significant with respect to stage to those found to be significant for survival. Second, we contrasted the matrisome genes and GO terms significant in each cohort with those in other cohorts, identifying similarities and differences between the gynecological cancers in terms of dysregulated and inferentially significant genes as well as dysregulated groups of GO terms. We propose that the variety of approaches in our multi- faceted analysis pipeline ameliorates the weaknesses of each constituent approach, yields informative sets of significantly dysregulated matrisome genes and enriched GO terms, and produces metrics which allow for sophisticated analyses of the relationship between matrisome gene expression and pathology.

Although the similarities between gynecological cancers are often the focal point of bioinformatics studies (Berger *et al*., 2018; Lim *et al*., 2019; Parris, 2020), our analyses demonstrate that at the matrisome level the transcriptomic profiles of these diseases are highly distinct from one another. Additionally, while gynecological cancers have been grouped with breast cancer in large pan-cancer studies (Whitley *et al*., 2004; Schüler-Toprak, Seitz and Ortmann, 2017; Berger *et al*., 2018; Ritter *et al*., 2018), the matrisome expression signatures of gynecological cancers were highly distinct from breast cancers. In fact, when compared to non-gynecological cancers, the matrisome transcriptome of gynecological cancers correlated more closely with non-gynecological cancers than with each other. Overall, the transcriptional matrisome characteristics of the gynecological cancers examined were more different than they are similar.

It has been widely demonstrated that a large proportion of genes are differentially expressed between normal and cancer tissues. In our work, we observed that this is especially true at the matrisome level, and even more pronounced for core-matrisome genes, which are less tissue-specific than ECM-related genes (Naba *et al*., 2012). Specifically, ECM GO terms were significantly enriched among the differentially expressed genes and a machine learning model trained on the matrisome transcriptome could almost perfectly distinguish normal from tumor tissue within each cancer type. Furthermore, in each of the three gynecological cancers evaluated, 65-75% of collagens and proteoglycans and 60-65% of ECM glycoproteins present in the dataset were dysregulated. Taken together, these results suggests that for future work seeking to characterize different types of cancer, the matrisome could be the most informative focal point. However, as with most large transcriptomic studies, it must be noted that the data we used are observational and no direct causal links can be established between the genes, gene networks, or gene sets and disease progression or patient survival using our findings. Thus *in vitro* or *in vivo* mechanistic studies should be performed to follow up on these observations.

Evaluating both univariable and multivariable models revealed key distinctions between gynecological cancer cohorts. Examining stage significant DEMGs revealed that CESC was characterized by diffuse gene ensemble behavior – many genes with weaker inferential significance for tumor stage – whereas UCEC and UCS had smaller ensembles of key genes acting together and serving as proxies for other tightly correlated genes. For patient survival, both univariable and multivariable analysis yielded several genes of interest in CESC, while patient survival results in UCEC and UCS were not easily characterized by individual DEMG expression levels. In UCEC, only *WIF1* was significant using DEMG-filtered univariable survival analysis. In UCS, only *TNFSF14* (linked to T cell proliferation, tumor cell apoptosis, and angiogenic normalization) and *CST1* (an inhibitor of cysteine proteinase found in several body fluids) were found to be significant via DEMG-filtered univariable survival analysis. WGCNA helped to fill in the gaps left by univariable and multivariable approaches in finding DEMGs significantly related to cancer stage, yielding several modules in each cohort which were significantly related to cancer stage. Within each gynecological cancer cohort, network modules yielded distinct GO terms. Taken together, these results indicate that analyses which seek to characterize cancer progression and patient survival should use both univariable and multivariable models along with network analysis, otherwise important gene markers will likely be overlooked.

Across FIGO stage significant DEMGs, differences between the cancers were observed, but many of the same downstream processes were impacted across cohorts. Results in CESC skewed towards immune regulation, underscored by widespread representation of cytokines and chemokines, likely due to the fact that over 90% of cervical cancer cases in the US are caused by HPV (Castellsagué, 2008; Saraiya *et al*., 2015; Centers for Disease Control, 2021). In contrast, UCEC and UCS results were biased toward ECM constituents and stromal signaling factors, without the same level of emphasis on immune regulation. All three cohorts had stage significant DEMGs linked to stromal signaling, cell growth, and structural ECM. The most prevalent stage significant DEMGs in all cohorts were proteases and protease inhibitors, collagen-containing ECM components, cytokines, immunomodulatory proteins, growth factor signaling proteins, and members of the semaphoring and plexin receptor-ligand families. Overall, dysregulation of ECM remodeling was common among all three gynecological cancer cohorts. In CESC and UCS, Cystatin E (*CST6*), which has been shown to contribute to cervical tumor growth and hyperactivation of cathepsin L when downregulated (Veena *et al*., 2008), was a stage significant DEMG. Proteases responsible for regulating core ECM components such as elastin, collagen, and fibronectin were also highly FIGO stage significant across cohorts. Specifically, MMPs were highly significant in all cohorts, cathepsins were significant in both CESC and UCEC, and the chymotrypsin-like elastase family was significant in UCS. While MMPs have previously been associated with FIGO stage in CESC via histology (Liu *et al*., 2018), this is the first work to demonstrate *in silico* that MMP expression is associated with FIGO stage in endometrial cancer and uterine sarcoma. Given that protease activity is critical for ECM remodeling and tumor cell invasion and migration (Winkler *et al*., 2020), it is unsurprising that we observed a strong presence of such enzymes among FIGO stage significant DEMGs in all cohorts. Numerous genes in the *ADAMTS* family, which coordinate pathophysiological ECM remodeling and regulate processes like inflammation and angiogenesis (Jones and Riley, 2005), were stage significant DEMGs in each of the three cancers. These genes target the members of the Von Willebrand Factor (VWF) family (Jones and Riley, 2005; South and Lane, 2018), which were also stage significant DEMGs in each cancer. The presence of both ADAMTS and VWF family proteins suggests a shared connection between FIGO stage and the formation of provisional ECM, likely as a means of promoting angiogenesis to support/sustain tumor growth (Feng *et al*., 2013). These results indicate that matrisome dysregulation related to tumor progression has common manifestations between cancer types, but significant distinctions are also apparent.

With respect to survival significant DEMGs, fewer similarities were observed between cancer cohorts. In CESC, many of the survival significant DEMGs were involved in immune modulation and regulation of immune cell binding. This builds upon previous work that demonstrates that an increase in tumor associated macrophages is associated with poor prognosis in cervical cancer patients (Chen *et al*., 2017). With respect to all cohorts, GO terms enriched among survival significant DEMGs were associated with processes which allow the cancer to evade the immune response and propagate in non-native tissues. While MMPs were shown to be significant among survival significant DEMGs in CESC, we did not identify any MMPs as predictive of survival in either UCS or UCEC. These results are similar to a separate study that evaluated which genes where predictive of patient survival in ovarian cancer (Vos *et al*., 2016). Overall, given the small number of death events in the data set (CESC: 66, UCEC: 24, UCS: 27), statistical power was limited, and survival significant genes were more difficult to identify than stage significant genes. As more survival data become available, future studies could potentially identify additional matrisome genes that are survival significant.

Generally, notable similarities and distinctions were observed among stage significant DEMGs between the three gynecological cancer cohorts, while survival significant DEMGs were largely different between the cohorts. Additionally, stage significant DEMGs among all three gynecological cancers contained more depth and breadth of gene family representation than survival significant DEMGs. Again, the small number of survival significant DEMGs is likely due to the small number of patient death events in the data and the impact of this on statistical power. One key finding in all three cohorts was the presence of FIGO stage and survival significant DEMGs involved in the WNT signaling pathway. WNT signaling is highly activated in all cancers (Zhan, Rindtorff and Boutros, 2017), and is modulated by changes in the ECM (Astudillo, 2020). In UCEC and CESC specifically, it has been associated with increased aggressiveness and cancer cell proliferation (Williams *et al*., 2017; Yang *et al*., 2018). In CESC, we observed that metalloproteinases such as SERPINs and MMPs, along with ECM assembly enzymes such as matrilins and lysyl oxidase (LOX)-family members, all of which are induced by WNT signaling (Wu, Crampton and Hughes, 2007; Li *et al*., 2018; Routledge and Scholpp, 2019), were prevalent among both stage and survival significant DEMGs. This is intuitive, as proteases promote ECM remodeling, tumorigenesis, and metastasis (Winkler *et al*., 2020). MMP-9 activity has been shown to be predictive of survival in cervical cancer patients, and our data further support this observation while also demonstrating that MMP-9 expression can be predictive of CESC FIGO stage (Li *et al*., 2012). Overall, stage and survival significant DEMGs in CESC were closely tied to ECM remodeling and immune signaling. UCEC stage and survival significant DEMGs consisted largely of genes involved in WNT signaling, which is often dysregulated in endometrial cancers (Fatima *et al*., 2021). In UCEC, we found that stage significant DEMGs were often within the WNT signaling pathway itself, whereas survival significant DEMGs were often upstream regulators of WNT signaling. There is a large body of evidence ascribing a role for WNT family signaling and dysregulation of the planar cell polarity pathway in oncogenic transformations through altered proliferation and pro-epithelial-to-mesenchymal transition behavior (VanderVorst *et al*., 2018). In low-grade endometrial cancers specifically, loss of epithelial apical polarity appears to drive proliferation and cancer cell migration (Williams *et al*., 2017). In UCS, stage and survival significant DEMG sets each shared an overrepresentation of the cystatin family of cysteine inhibitors as well as the mucin family, both of which are regulated by the WNT pathway (Lee *et al*., 2015; Pai *et al*., 2016). While little research exists on the role of these proteins in UCS, mucins appear to be elevated in ovarian carcinomas and are currently being investigated as a diagnostic candidate (Bonifácio, 2020). With regard to genes directly involved in WNT signaling, UCS may be characterized less by dysregulation of these components due to its mesenchymal tissue origin, meaning there is a reduced need for the tumor tissue to undergo epithelial-mesenchymal transition. Taken together, our results confirm the importance of the WNT signaling pathway across all three gynecological cancer cohorts investigated. Furthermore, our results identified key similarities and differences between stage and survival significant matrisome dysregulation, and support and expand upon previous findings in the literature. Though our results point to many contrasts between the cancer cohorts, it should be noted that the unified dataset used in this study, while corrected for batch effects, likely retains some variance attributable to combining data from different data sources.

Our findings have immediate relevance to the fields of cancer tissue engineering. The role of regulators of ECM turnover/remodeling versus physical constituents in each cancer should be further investigated to determine influence on tumor progression. Engineered tissue systems often utilize hydrogels coated with a single ECM protein (e.g., collagen I, fibronectin) or synthetic polymers possessing adhesion peptides, but we show in this work that the key drivers of tumor progression and patient mortality are not the adhesion peptides or structural proteins of the ECM, but rather the proteins which dynamically regulate ECM turnover. Though there was a notable amount of overlap between each of the three cancer cohorts in terms of stage and survival DEMGs, the matrisome gene signatures unique to each cohort could be utilized to inform biomaterial constructs aimed at studying tumor progression or studying cancer cell behavior in environments deterministic of low survival rates. For example, extracellular proteases were frequently found among stage significant DEMGs in each of the three gynecological cancers. Therefore, incorporating protease cleavable sites into a synthetic material construct, such as MMP-cleavable peptide sequences, similar to the work done by Valdez *et al*. (Valdez *et al*., 2017), could be useful for studying ECM turnover and cancer cell invasion. Likewise, the inclusion of heparin or cleavable heparin-binding sites that sequester soluble factors such as TGF-β would enable controlled release during matrix remodeling.

Techniques for controlled delivery of heparin-binding growth factors during tissue regeneration are discussed at length by Joung *et al*. (Joung, Bae and Park, 2008). Information gleaned from our study could also be used to create novel material constructs for each cancer comprised of distinct features specific to either tumor stage or patient survival. We previously noted that DEMGs significant for patient survival in CESC favored regulation of immune signaling and recruitment whereas those significant in UCS and UCEC were biased towards stromal signaling factors. This information could be applied to engineering 3D *in vitro* models with spatially distinct regions of adhesive peptides and tethered growth factors. *In vitro* models of CESC could incorporate gradients of various, pro-tumorigenic cytokines identified in our results, such as IL- 17, which is associated with tumor growth (Zhao *et al*., 2019). Similarly, we noted an overrepresentation of laminins and various proteoglycan-associated proteins among stage significant DEMGs in UCS and UCEC. This overrepresentation may imply a role for incorporating DEMGs associated with a pro-invasion basement membrane into a tissue model system. Finally, the presence of collagens and various collagen processing components in both stage and survival significant DEMG sets across all cancers further underscores the role for collagen organization and density as deterministic of cancer aggressiveness (Acerbi *et al*., 2015). Taken together, our results support the consideration of biophysical cues such as stiffness, controlled release of soluble factors, and inclusion of protease cleavable sites when designing *in vitro* constructs to study gynecological cancers. Furthermore, our results identify key pathways that should be considered when designing disease-specific *in vitro* models.

The results of this work can provide an atlas for further matrisome gene expression research in the context of the gynecological cancers studied. While these diseases are wide-spread and deadly for women worldwide, concerted efforts to broadly characterize their similarities and distinctions, especially at the matrisome level, are not prevalent. Our research contributes to a better understanding of the matrisome pathways that are dysregulated in gynecological cancers. These pathways can be further investigated with clinical samples, *in vivo* models, or 3D *in vitro* models. Furthermore, our findings identify disease-specific dysregulated pathways that can be investigated as potential therapeutic targets.

## Methods

### Data sources and preprocessing

All data preprocessing was done using the *R* (R Core Team, 2020) and *Python* programming languages. The terms “tumor” or “tumor sample” refer specifically to primary tumor samples from the given dataset.

### Unified dataset

The un-normalized unified RNA-Seq dataset of The Cancer Genome Atlas (TCGA) gynecological tumor samples and Genotype Tissue Expression (GTEx) database samples created by Wang *et al*. was accessed and downloaded from the Figshare scientific data sharing website (https://figshare.com/articles/dataset/Data_record_1/5330539) (Q. Wang *et al*., 2018) on 7/20/2020. TCGA is a repository of several petabytes of publicly available cancer data, hosted on the Genomic Data Commons portal (Tomczak, Czerwińska and Wiznerowicz, 2015).

While large numbers of tumor samples are available from TCGA, normal (healthy) samples are lacking. GTEx is a public repository with a wealth of gene expression data generated from normal tissue samples (Aguet *et al*., 2017), making it an ideal source for additional data to pair with TCGA. The unified dataset created by Wang *et al*. combines the TCGA and GTEx data (for cancer types with both tumor and normal samples present in the TCGA RNA-seq database) and processes them uniformly, removing batch effects. The raw data were used for differential gene expression (DGE) analysis. For other analyses of these data, we employed the *varianceStabilizingTransofrmation* function from the *DESeq2* (Love, Huber and Anders, 2014) package to reduce the amount of heteroskedasticity in the gene expression count data. We refer to these data as the variance-stabilizing transformed (VST) unified data. Unfortunately, due to the lack of normal ovarian tissue samples present in the TCGA dataset, the ovarian cohort was not included by Wang *et al*. and is used by us only for the investigation of tumor heterogeneity in our analyses (since it cannot be subjected to DGE analysis without any normal tissue samples).

### TCGA count & clinical data

TCGA *HTSeq* count data (NCI Genomic Data Commons, 2021) and the corresponding clinical data (e.g., demographics, FIGO stage, survival) were downloaded from TCGA via the *TCGABiolinks* package (Colaprico *et al*., 2016) on 2020/09/15. The gene expression count data (normalized using the *varianceStabilizingTransofrmation* function from *DESeq2*) were used for our heterogeneity analysis, while the clinical data were matched with corresponding data from the unified dataset for use in our other analyses. For the heterogeneity analysis, we downloaded data corresponding to breast invasive carcinoma (BRCA), cervical squamous cell carcinoma and endocervical adenocarcinoma (CESC), colon adenocarcinoma (COAD), ovarian serous cystadenocarcinoma (OV), prostate adenocarcinoma (PRAD), testicular germ cell tumors (TGCT), uterine corpus endometrial carcinoma (UCEC), uterine carcinosarcoma (UCS).

### The matrisome database

The human matrisome database was constructed by Naba *et al*. using a combination bioinformatic protein domain sweeps and manual curation (Naba *et al*., 2012). The human matrisome database was retrieved from the online repository of The Matrisome Project (http://matrisomeproject.mit.edu/other-resources/human-matrisome) on 7/21/2020. We filtered out the “retired” genes in this database, yielding a master list of 1027 genes, 1008 of which were present in the unified dataset used in our analyses. The 1027-gene master list consisted of core matrisome (*n* = 274) and matrisome-associated (*n* = 753) genes. Matrisome categories included: collagens, ECM glycoproteins, ECM regulators, ECM-affiliated proteins, proteoglycans, and secreted factors (**Table S7**). Of these, 270 core matrisome and 738 matrisome-associated genes (1,008 total) were present in our dataset (**Table S7**).

### Correlative hierarchical clustering

To visualize the overall heterogeneity of matrisome expression among gynecological cancers, correlative hierarchical clustering was performed on select cancer types in TCGA. This was achieved using the *pheatmap* function from the *pheatmap* package in *R* (Kolde, 2019). We selected 25 representative samples from each cancer cohort by first computing centroids for each cohort (gene-wise *median* expression values) and selected the 25 tumor samples which were closest (as measured by L^1^ distance) to these centroids. This procedure can be summarized by the expression

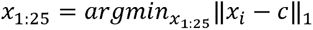

Where *x*_125_ denotes the 25 samples with minimal L^1^ distance from the cohort centroid, *c*. This sample pre-selection was done to ensure that the samples being compared via clustering were as representative of the cancer types to which they belonged as possible. Clustering was performed using the *hclust* function from *stats* package in *R* (R Core Team, 2020). The distance metric used was 1 - *r_ij_*, where *r_ij_* is the correlation coefficient for the th and *i*th and *j*th genes (or samples). Pearson correlation was used for gene-wise correlation, while Spearman correlation was used for sample-wise correlation. This clustering was done on the TCGA *HTSeq* count data, normalized using the *varianceStabilizingTransformation* function from *DESeq2*.

### Dimensionality reduction

Dimensionality reduction was performed for the TCGA *HTSeq* count data, normalized using the *varianceStabilizingTransformation* function from *DESeq2*. PCA was performed using the *prcomp* function from *R*’s *stats* package. UMAP (McInnes, Healy and Melville, 2020) dimensionality reduction was performed using the *umap* function from the *umap* package in *R* (Konopka, 2020). While UMAP analysis is predominantly used in single cell RNA-Seq data analysis (Becht *et al*., 2019), we applied it to bulk RNA-Seq data to visually stratify cancer type sub-populations in 2-dimensional plots.

### Tumor versus normal tissue stratification by machine learning

Elastic net logistic regression models (Algamal and Lee, 2015) were trained on the VST unified matrisome expression data for each gynecological cancer cohort. These models were trained to classify, in each cohort, observations as tumor samples or normal samples to demonstrate the power of matrisome expression profiles in characterizing cancers. Elastic net regression utilizes the objective function

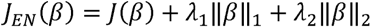

where *J*(*β*) is a simpler loss function and the parameters *λ*_1_ and *λ*_2_ control the proportion of L^1^ (lasso) and L^2^ (ridge) regression penalization. The performance of these models was measured using balanced classification accuracy (Brodersen *et al*., 2010). This scoring method utilizes observation weights defined according to

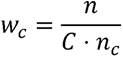

where *w_C_* is the weight assigned to observations from class, lll is the total number of observations, is the total number of classes (factor levels of the response), and *C* is the total number of classes (factor levels of the response), and *n_c_* is the number of observations in class *c* (Pedregosa *et al*., 2011). The weights sum to 1 and ensure that the sum of the weighted observations of each class are the same, penalizing models which only perform well on the most common class(es) in the data set. These models were trained using 5-fold cross validation and hyperparameters were optimized using sequential model-based optimization (Hutter, Hoos and Leyton-Brown, 2011). The scikit-learn implementation of elastic net logistic regression was used (Pedregosa *et al*., 2011), and sequential model-based optimization was performed using the *gp_minimize* function in scikit-optimize (Head *et al*., 2018).

### Differential Gene Expression analysis

Differential gene expression (DGE) analysis was conducted on the full unified data set using the *DESeq2* package in *R* (Love, Huber and Anders, 2014). All genes with expression counts of 0 in more than one third of samples were considered to be lowly expressed genes and filtered out prior to performing the differential gene expression analysis. Using the recommended two-step procedure (Love, Anders and Huber, 2021), the *DESeq* function was called with default arguments, followed by the *results* function with *alpha* set to the significance threshold, *α* = 0.05, and sample condition modeled as tumor versus normal. Genes were deemed differentially expressed based on two commonly used criteria: 1) *q*-value less than the significance threshold (*α* = 0.05) and 2) absolute log fold-change of 1 (McCarthy and Smyth, 2009). While the Benjamini-Hoschberg option was used when calling the *DESeq* function, we determined differential expression based on results from the more sophisticated *qvalue* function from the *WGCNA* package in R (Langfelder and Horvath, 2008). In the case of UCS and UCEC, the same normal uterine tissue samples, were used for determining differential expression.

### Univariable statistical analyses

Several large statistical analyses were performed on the unified gene expression data (tumor samples only). Aside from the pairwise FIGO differential gene expression analysis, which utilized count data for all genes, the VST expression data for the matrisome genes was used. Once again, *q*-values were computed by adjusting -values using the *WGCNA* package’s *qvalue* function. For analyses involving FIGO stage, all sub-stages were condensed into stages I-IV (for example, Stage IA and Stage IB would be replaced by Stage I) and observations with missing FIGO data were removed. For analyses involving censored survival time, samples with missing survival time or status (alive or deceased) or survival time of 0 days were filtered out.

### Pairwise FIGO differential expression analysis

Differential expression analysis was performed pairwise between tumor samples from all different FIGO stages. The following contrasts were used (numbers correspond to FIGO stages): 2 vs. 1, 3 vs. 1, 4 vs. 1, 3 vs. 2, 4 vs. 2, and 4 vs. 3. Results were combined and filtered to include only matrisome genes. Significance was determined by Benjamini-Hochberg adjusted *p*-values (*p_adj_* < 0.01) and absolute log fold-change of 1 or more. A matrisome gene was required to meet both the adjusted -value and log fold-change threshold in at least one contrast to be considered significant. Our motivation for using stricter false discovery rate controls than in the earlier tumor versus healthy tissue differential expression analysis was to create more parsimonious lists with lower risk of type I error rate. This is because, 1) the analysis was performed on all genes and results were filtered to include only matrisome genes, creating the potential for a mismatch in intended and achieved FDR among the target population and 2) the individual genes discovered in this analysis would be used for detailed downstream analysis and high confidence in the results was desired. These analyses were completed using the *DESeq* function with parameter *alpha* set to 0.01, the intended significance threshold.

### Point-biserial correlation

Point-biserial correlation is mathematically equivalent to the Pearson correlation between a continuous and dichotomous variable (Kornbrot, 2005). The FIGO stage of each sample was transformed using a one hot encoding and gene-wise point-biserial correlations were computed for each FIGO stage. Genes were deemed significant if their Student asymptotic *q*-values (Langfelder and Horvath, 2008) were below the significance threshold *q* < 0.05) for at least one FIGO stage. Student asymptotic *p*-values were computed using the *corPvalueStudent* function from the *WGCNA* package and then adjusted.

### Censored time screening

The *standardScreeningCensoredTime* function from the WGCNA package was used to test whether any matrisome genes were significantly related to patient survival (log-rank test tl < 0.05). The authors describe this method as fitting gene-wise univariable Cox PH models and performing log-rank tests (Langfelder and Horvath, 2008).

### Univariable Kaplan-Meier and Cox PH

Each matrisome gene was related to survival via the union of Kaplan-Meier and univariable Cox PH models (Kaplan and Meier, 1958; Cox, 1972). For Kaplan-Meier analysis, a gene expression cutoff first had to be set for each gene, so that they could be divided into high and low expression groups. This cutoff was found by fitting a two component Gaussian mixture model to the VST expression data each gene and setting the cutoff to be equal to the expression value at which the two Gaussian densities most closely overlapped, similar to the method described by Budczies *et al*. (Budczies *et al*., 2012). Gene-wise Kaplan-Meier modeling was then performed on these stratified expression groups. Gene-wise Cox PH models were fit to the unstratified VST expression data. Genes were classified as significant based on their Kaplan-Meier *q*-values (Chi-squared test *q* < 0.05) or Cox PH *q* -values (likelihood ratio test *q* < 0.05). While use of gene-wise univariable Cox PH in this method may seem redundant with respect to the censored time screening, the two methods employed different hypothesis tests and did not have perfectly overlapping results, so both were included.

### Machine learning-based feature selection

L^1^ penalized multivariable machine learning models were fit to the VST unified matrisome data for tumor samples in each gynecological cancer cohort. The purpose of these models was to identify relationships between matrisome genes and both FIGO stage and patient survival. The same pre-filtering that was used for the univariable statistical FIGO and survival analyses was repeated here. In both cases, an L^1^ penalized model was fit to the data using the *glmnet* package in *R* (Friedman, Hastie and Tibshirani, 2010; Simon *et al*., 2011). The models were optimized for parsimony, which is done by tuning the value of *λ* in the L^1^ penalty objective function

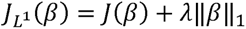

where *J*(*β*) is some simpler loss function (misclassification rate, for example) and ||*β*|| is the L^1^ norm of the model parameters. FIGO and survival models were fit in each gynecological cohort using cross validation and a 1-dimensional grid search over values of ’:f. The performances of the worst FIGO model and survival model were used as caps for the models in the other cohorts. This procedure consisted of identifying (for both FIGO models and survival models) the best score within each cohort and then selecting the worst of these (i.e., the worst-of-the-best); this score was then used to select the first model which achieved a comparable (to 3 decimal places) score in each of the other cohorts (**Figure S4**). This was done for two reasons; first, since model results would be compared between cancer cohorts, it was desirable to compare results from models of similar predictive power. Second, the lowest performing models were also the least parsimonious, so employing this restriction had the effect of bounding how parsimonious any model could be – to avoid, for example, comparing models which selected 300 features to those which selected 3. Values of ’:f were searched in increasing order during model fitting, where 1000 evenly spaced values of ’:f were searched on a log scale in the range [G^-^, G^O^] (**Figure S4**).

### FIGO predictive multinomial regression

Four-class multinomial regression was used to select genes which were related to FIGO stage, since FIGO stage classification was a four-level classification problem. The poorest performance was observed in the CESC cohort (**Table S8**). The multinomial regressors were fit and assessed using 5-fold cross validation (10-fold cross validation was counter-indicated because 10 observations were not present for every level of FIGO stage in every cohort) by using the *cv.glmnet* function in the *glmnet* package with arguments *family* set to “multinomial”, *type.multinomial* set to “grouped” to ensure coefficients at the same indices (for the same genes) would be retained for each level of the response, and *type.measure* set to “class” meaning classification error was used. Balanced class-weighting (as in the tumor/normal stratification models) was used to ensure models did not favor performance in any FIGO stage. This balanced class-weighting was critical since FIGO stages I-IV were highly imbalanced.

### Survival predictive Cox proportional hazards

Multivariable Cox PH regression was used to select genes which were related to survival. The poorest performance was observed in UCS (**Table S8**), which was unsurprising since this was the smallest cohort. The Cox PH regressors were fit and assessed using 10-fold cross validation using the *cv.glmnet* function in the *glmnet* package with arguments *family* set to “cox” and *type.measure* set to “C” for Harrel’s concordance measure (Steck *et al*., 2007).

### Weighted gene correlation network analysis (WGCNA)

WGCNA was performed according to the instructions provided by the package authors (Langfelder and Horvath, 2016). First, a topological overlap matrix was constructed over the matrisome genes using a soft power which yielded a sufficient scale free topological overlap metric (TOM) (Zhang and Horvath, 2005). This matrix of TOMs represents the gene-wise similarity of expression values for each matrisome gene, with more similar genes having larger topological overlap values. This matrix was then used to perform hierarchical clustering, where modules were merged if they achieved a correlative distance threshold of 0.25 (i.e., module correlation of 0.75). Modules were then related to two traits: *censored survival time* and FIGO stage. Module-trait significance was determined by correlating the module *eigengenes* with the traits of interest. Matrisome genes were determined to be significant if 1) they belonged to a module with a significant FIGO correlation for at least one FIGO stage (Student asymptotic tl < 0.05) and 2) their module membership was significant (Student asymptotic <). All WGCNA was performed using *R* code and functions from the *WGCNA* package in *R* (Langfelder and Horvath, 2008).

### Enrichment analysis

Gene set and pathway enrichment analysis were performed using the *R* package, *clusterProfiler* (Wu *et al*., 2021). For gene set (functional) enrichment analysis, the function *enrichGO* was used to find enriched gene ontologies (GO) among significant genes (Gene Ontology Consortium, 2004). For pathway enrichment analysis, the function *enrichKEGG* was used to find KEGG pathways which were enriched among significant genes (Kanehisa and Goto, 2000). The significance of each enriched gene function or pathway was determined based on the reported *q*-value (*q* < 0.05).

### Qualitative analysis of FIGO and Survival-significant gene lists

Identification and overall relevancy of overrepresented gene families were identified using STRING database queries within Cytoscape (v3.8.0) software. To compare both within and between FIGO stage/survival significant genes for each cancer, lists of transcripts were compared on a gene-by-gene (e.g., *MMP3)* and family-by-family (e.g., Mucins) basis.

## Table Captions

*Table 1. Differential gene expression summary for tumor versus healthy tissue in each gynecological cancer cohort*.

Breakdown of DEGs & DEMGs by cancer cohort (CESC, UCEC, and UCS). Here, DE means differentially expressed and % DE means percent of all/matrisome genes found to be differentially expressed. Sample sizes CESC (*n_normal_* = 13, *n_tumor_* =259), UCEC (*n_normal_* = 105, *n_tumor_* = 141), and UCS (*n_normal_* = 105, *n_tumor_* = 47).

## References

Acerbi, I. et al. (2015) ‘Human breast cancer invasion and aggression correlates with ECM stiffening and immune cell infiltration’, Integrative Biology: Quantitative Biosciences from Nano to Macro, 7(10), pp. 1120–1134. doi:10.1039/c5ib00040h.

Aguet, F. et al. (2017) ‘Genetic effects on gene expression across human tissues’, Nature, 550(7675), pp. 204–213. doi:10.1038/nature24277.

Algamal, Z.Y. and Lee, M.H. (2015) ‘Applying Penalized Binary Logistic Regression with Correlation Based Elastic Net for Variables Selection’, Journal of Modern Applied Statistical Methods, 14(1), pp. 168–179. doi:10.22237/jmasm/1430453640.

Apte, S.S. (2009) ‘A Disintegrin-like and Metalloprotease (Reprolysin-type) with Thrombospondin Type 1 Motif (ADAMTS) Superfamily: Functions and Mechanisms’, The Journal of Biological Chemistry, 284(46), pp. 31493–31497. doi:10.1074/jbc.R109.052340.

Arbyn, M. et al. (2020) ‘Estimates of incidence and mortality of cervical cancer in 2018: a worldwide analysis’, The Lancet Global Health, 8(2), pp. e191–e203. doi:10.1016/S2214-109X(19)30482-6.

Ashburner, M. et al. (2000) ‘Gene Ontology: tool for the unification of biology’, Nature genetics, 25(1), pp. 25–29. doi:10.1038/75556.

Astudillo, P. (2020) ‘Extracellular matrix stiffness and Wnt/β-catenin signaling in physiology and disease’, Biochemical Society Transactions, 48(3), pp. 1187–1198. doi:10.1042/BST20200026.

Becht, E. et al. (2019) ‘Dimensionality reduction for visualizing single-cell data using UMAP’, Nature Biotechnology, 37(1), pp. 38–44. doi:10.1038/nbt.4314.

Berger, A.C. et al. (2018) ‘A Comprehensive Pan-Cancer Molecular Study of Gynecologic and Breast Cancers’, Cancer Cell, 33(4), pp. 690–705.e9. doi:10.1016/j.ccell.2018.03.014.

Bonifácio, V.D.B. (2020) ‘Ovarian Cancer Biomarkers: Moving Forward in Early Detection’, Advances in Experimental Medicine and Biology, 1219, pp. 355–363. doi:10.1007/978-3-030-34025-4_18.

Braun, R. et al. (2013) ‘Discovery Analysis of TCGA Data Reveals Association between Germline Genotype and Survival in Ovarian Cancer Patients’, PLOS ONE, 8(3), p. e55037. doi:10.1371/journal.pone.0055037.

Brodersen, K.H. et al. (2010) ‘The Balanced Accuracy and Its Posterior Distribution’, in 2010 20th International Conference on Pattern Recognition. 2010 20th International Conference on Pattern Recognition, pp. 3121–3124. doi:10.1109/ICPR.2010.764.

Bruchim, I., Sarfstein, R. and Werner, H. (2014) ‘The IGF Hormonal Network in Endometrial Cancer: Functions, Regulation, and Targeting Approaches’, Frontiers in Endocrinology, 5. doi:10.3389/fendo.2014.00076.

Budczies, J. et al. (2012) ‘Cutoff Finder: A Comprehensive and Straightforward Web Application Enabling Rapid Biomarker Cutoff Optimization’, PLoS ONE, 7(12). doi:10.1371/journal.pone.0051862.

Castellsagué, X. (2008) ‘Natural history and epidemiology of HPV infection and cervical cancer’, Gynecologic Oncology, 110(3, Supplement 2), pp. S4–S7. doi:10.1016/j.ygyno.2008.07.045.

Centers for Disease Control (2020) What Is Gynecologic Cancer? Available at: https://www.cdc.gov/cancer/gynecologic/basic_info/what-is-gynecologic-cancer.htm (Accessed: 26 April 2021).

Centers for Disease Control (2021) How Many Cancers Are Linked with HPV Each Year? Available at: https://www.cdc.gov/cancer/hpv/statistics/cases.htm (Accessed: 18 August 2021).

Cestarelli, V. et al. (2016) ‘CAMUR: Knowledge extraction from RNA-seq cancer data through equivalent classification rules’, Bioinformatics, 32(5), pp. 697–704. doi:10.1093/bioinformatics/btv635.

Chen, X.-J. et al. (2017) ‘Clinical Significance of CD163+ and CD68+ Tumor-associated Macrophages in High-risk HPV-related Cervical Cancer’, Journal of Cancer, 8(18), pp. 3868– 3875. doi:10.7150/jca.21444.

Cherniack, A.D. et al. (2017) ‘Integrated Molecular Characterization of Uterine Carcinosarcoma’, Cancer Cell, 31(3), pp. 411–423. doi:10.1016/j.ccell.2017.02.010.

Clayton, E.A. et al. (2020) ‘Leveraging TCGA gene expression data to build predictive models for cancer drug response’, BMC Bioinformatics, 21(14), p. 364. doi:10.1186/s12859-020-03690-4.

Cline, M.S. et al. (2013) ‘Exploring TCGA Pan-Cancer Data at the UCSC Cancer Genomics Browser’, Scientific Reports, 3(1), p. 2652. doi:10.1038/srep02652.

Colaprico, A. et al. (2016) ‘TCGAbiolinks: an R/Bioconductor package for integrative analysis of TCGA data’, Nucleic Acids Research, 44(8), pp. e71–e71. doi:10.1093/nar/gkv1507.

Cox, D.R. (1972) ‘Regression Models and Life-Tables’, Journal of the Royal Statistical Society: Series B (Methodological), 34(2), pp. 187–202. doi:https://doi.org/10.1111/j.2517-6161.1972.tb00899.x.

Dai, X. et al. (2019) ‘Identifying Interaction Clusters for MiRNA and MRNA Pairs in TCGA Network’, Genes, 10(9), p. 702. doi:10.3390/genes10090702.

Dellinger, T.H. et al. (2016) ‘L1CAM is an independent predictor of poor survival in endometrial cancer – an analysis of The Cancer Genome Atlas (TCGA)’, Gynecologic oncology, 141(2), pp. 336–340. doi:10.1016/j.ygyno.2016.02.003.

Facciabene, A. et al. (2011) ‘Tumour hypoxia promotes tolerance and angiogenesis via CCL28 and Treg cells’, Nature, 475(7355), pp. 226–230. doi:10.1038/nature10169.

Fatima, I. et al. (2021) ‘Targeting Wnt Signaling in Endometrial Cancer’, Cancers, 13(10), p. 2351. doi:10.3390/cancers13102351.

Felix, A.S. and Brinton, L.A. (2018) ‘Cancer Progress and Priorities: Uterine Cancer’, Cancer epidemiology, biomarkers & prevention : a publication of the American Association for Cancer Research, cosponsored by the American Society of Preventive Oncology, 27(9), pp. 985–994. doi:10.1158/1055-9965.EPI-18-0264.

Feng, X. et al. (2013) ‘Fibrin and collagen differentially but synergistically regulate sprout angiogenesis of human dermal microvascular endothelial cells in 3-dimensional matrix’, International Journal of Cell Biology, 2013, p. 231279. doi:10.1155/2013/231279.

Frantz, C., Stewart, K.M. and Weaver, V.M. (2010) ‘The extracellular matrix at a glance’, Journal of Cell Science, 123(24), pp. 4195–4200. doi:10.1242/jcs.023820.

Freeman, S.J. et al. (2012) ‘The Revised FIGO Staging System for Uterine Malignancies: Implications for MR Imaging’, RadioGraphics, 32(6), pp. 1805–1827. doi:10.1148/rg.326125519.

Friedman, J., Hastie, T. and Tibshirani, R. (2010) ‘Regularization Paths for Generalized Linear Models via Coordinate Descent’, Journal of Statistical Software, 33(1). doi:10.18637/jss.v033.i01.

Funston, G. et al. (2018) ‘Recognizing Gynecological Cancer in Primary Care: Risk Factors, Red Flags, and Referrals’, Advances in Therapy, 35(4), pp. 577–589. doi:10.1007/s12325-018-0683-3.

Gene Ontology Consortium (2004) ‘The Gene Ontology (GO) database and informatics resource’, Nucleic Acids Research, 32(suppl_1), pp. D258–D261. doi:10.1093/nar/gkh036.

Goeman, J.J. (2010) ‘L1 Penalized Estimation in the Cox Proportional Hazards Model’, Biometrical Journal, 52(1), pp. 70–84. doi:https://doi.org/10.1002/bimj.200900028.

Head, T. et al. (2018) scikit-optimize. Zenodo. doi:10.5281/zenodo.1207017.

Henley, S.J. et al. (2018) ‘Uterine Cancer Incidence and Mortality — United States, 1999–2016’, Morbidity and Mortality Weekly Report, 67(48), pp. 1333–1338. doi:10.15585/mmwr.mm6748a1.

Hutter, F., Hoos, H.H. and Leyton-Brown, K. (2011) ‘Sequential Model-Based Optimization for General Algorithm Configuration’, in Coello, C.A.C. (ed.) *Learning and Intelligent Optimization*. Berlin, Heidelberg: Springer (Lecture Notes in Computer Science), pp. 507–523. doi:10.1007/978-3-642-25566-3_40.

Hynes, R.O. and Naba, A. (2012) ‘Overview of the matrisome--an inventory of extracellular matrix constituents and functions’, Cold Spring Harbor Perspectives in Biology, 4(1), p. a004903. doi:10.1101/cshperspect.a004903.

Jha, A. et al. (2017) ‘Towards precision medicine: discovering novel gynecological cancer biomarkers and pathways using linked data’, Journal of Biomedical Semantics, 8(1), p. 40. doi:10.1186/s13326-017-0146-9.

Jones, G.C. and Riley, G.P. (2005) ‘ADAMTS proteinases: a multi-domain, multi-functional family with roles in extracellular matrix turnover and arthritis’, Arthritis Research & Therapy, 7(4), p. 160. doi:10.1186/ar1783.

Joung, Y.K., Bae, J.W. and Park, K.D. (2008) ‘Controlled release of heparin-binding growth factors using heparin-containing particulate systems for tissue regeneration’, Expert Opinion on Drug Delivery, 5(11), pp. 1173–1184. doi:10.1517/17425240802431811.

Kanehisa, M. and Goto, S. (2000) ‘KEGG: Kyoto Encyclopedia of Genes and Genomes’, Nucleic Acids Research, 28(1), pp. 27–30. doi:10.1093/nar/28.1.27.

Kaplan, E.L. and Meier, P. (1958) ‘Nonparametric Estimation from Incomplete Observations’, Journal of the American Statistical Association, 53(282), pp. 457–481. doi:10.1080/01621459.1958.10501452.

Khoury, J.D. et al. (2013) ‘Landscape of DNA Virus Associations across Human Malignant Cancers: Analysis of 3,775 Cases Using RNA-Seq’, Journal of Virology, 87(16), pp. 8916–8926. doi:10.1128/JVI.00340-13.

Kolde, R. (2019) pheatmap: Pretty Heatmaps. Available at: https://CRAN.R-project.org/package=pheatmap (Accessed: 16 February 2021).

Konopka, T. (2020) umap: Uniform Manifold Approximation and Projection. Available at: https://CRAN.R-project.org/package=umap (Accessed: 16 February 2021).

Kornbrot, D. (2005) ‘Point Biserial Correlation’, in Encyclopedia of Statistics in Behavioral Science. American Cancer Society. doi:10.1002/0470013192.bsa485.

Langfelder, P. and Horvath, S. (2008) ‘WGCNA: an R package for weighted correlation network analysis’, BMC Bioinformatics, 9(1), p. 559. doi:10.1186/1471-2105-9-559.

Langfelder, P. and Horvath, S. (2016) Tutorials for WGCNA R package, Tutorials for the WGCNA package. Available at: https://horvath.genetics.ucla.edu/html/CoexpressionNetwork/Rpackages/WGCNA/Tutorials/ (Accessed: 16 February 2021).

Langhans, S.A. (2018) ‘Three-Dimensional in Vitro Cell Culture Models in Drug Discovery and Drug Repositioning’, Frontiers in Pharmacology, 9. doi:10.3389/fphar.2018.00006.

Lee, H.-J. et al. (2015) ‘Structure-based Discovery of Novel Small Molecule Wnt Signaling Inhibitors by Targeting the Cysteine-rich Domain of Frizzled’, The Journal of Biological Chemistry, 290(51), pp. 30596–30606. doi:10.1074/jbc.M115.673202.

Li, H. et al. (2018) ‘LOXL1 regulates cell apoptosis and migration in human neuroglioma U87 and U251 cells via Wnt/β-catenin signaling’, International Journal of Clinical and Experimental Pathology, 11(4), pp. 2032–2037.

Li, Y. et al. (2012) ‘Matrix metalloproteinase-9 is a prognostic marker for patients with cervical cancer’, Medical Oncology (Northwood, London, England), 29(5), pp. 3394–3399. doi:10.1007/s12032-012-0283-z.

Lim, S.B. et al. (2019) ‘Pan-cancer analysis connects tumor matrisome to immune response’, npj Precision Oncology, 3(1), pp. 1–9. doi:10.1038/s41698-019-0087-0.

Liu, C. et al. (2018) ‘Clinical significance of matrix metalloproteinase-2 in endometrial cancer: A systematic review and meta-analysis’, Medicine, 97(29), p. e10994. doi:10.1097/MD.0000000000010994.

Lopez Vicchi, F. and Becu-Villalobos, D. (2017) ‘Prolactin: The Bright and the Dark Side’, Endocrinology, 158(6), pp. 1556–1559. doi:10.1210/en.2017-00184.

Love, M.I., Anders, S. and Huber, W. (2021) Analyzing RNA-seq data with DESeq2, Bioconductor. Available at: https://bioconductor.org/packages/release/bioc/vignettes/DESeq2/inst/doc/DESeq2.html (Accessed: 16 February 2021).

Love, M.I., Huber, W. and Anders, S. (2014) ‘Moderated estimation of fold change and dispersion for RNA-seq data with DESeq2’, Genome Biology, 15(12), p. 550. doi:10.1186/s13059-014-0550-8.

Massagué, J. (2008) ‘TGFβ in Cancer’, Cell, 134(2), pp. 215–230. doi:10.1016/j.cell.2008.07.001.

Mathur, S.P. et al. (2003) ‘Circulating levels of insulin-like growth factor-II and IGF-binding protein 3 in cervical cancer’, Gynecologic Oncology, 91(3), pp. 486–493. doi:10.1016/j.ygyno.2003.08.023.

Matsuo, K. et al. (2016) ‘Significance of histologic pattern of carcinoma and sarcoma components on survival outcomes of uterine carcinosarcoma’, Annals of Oncology, 27(7), pp. 1257–1266. doi:10.1093/annonc/mdw161.

McCarthy, D.J. and Smyth, G.K. (2009) ‘Testing significance relative to a fold-change threshold is a TREAT’, Bioinformatics, 25(6), pp. 765–771. doi:10.1093/bioinformatics/btp053.

McInnes, L., Healy, J. and Melville, J. (2020) ‘UMAP: Uniform Manifold Approximation and Projection for Dimension Reduction’, arXiv:1802.03426 [cs, stat] [Preprint]. Available at: http://arxiv.org/abs/1802.03426 (Accessed: 28 January 2021).

Mouw, J.K. et al. (2014) ‘Tissue mechanics modulate microRNA-dependent PTEN expression to regulate malignant progression’, Nature Medicine, 20(4), pp. 360–367. doi:10.1038/nm.3497.

Naba, A. et al. (2012) ‘The matrisome: in silico definition and in vivo characterization by proteomics of normal and tumor extracellular matrices’, Molecular & cellular proteomics: MCP, 11(4), p. M111.014647. doi:10.1074/mcp.M111.014647.

Naba, A., Hoersch, S. and Hynes, R.O. (2012) ‘Towards definition of an ECM parts list: An advance on GO categories’, Matrix biology :journal of the International Society for Matrix Biology, 31(0), pp. 371–372. doi:10.1016/j.matbio.2012.11.008.

NCI Genomic Data Commons (2021) GDC Docs. Available at: https://docs.gdc.cancer.gov/Encyclopedia/pages/HTSeq-Counts/ (Accessed: 16 February 2021).

Pai, P. et al. (2016) ‘Mucins and Wnt/β catenin signaling in gastrointestinal cancers: an unholy nexus’, Carcinogenesis, 37(3), pp. 223–232. doi:10.1093/carcin/bgw005.

Parris, T.Z. (2020) ‘Pan-cancer analyses of human nuclear receptors reveal transcriptome diversity and prognostic value across cancer types’, Scientific Reports, 10(1), p. 1873. doi:10.1038/s41598-020-58842-6.

Pedregosa, F. et al. (2011) ‘Scikit-learn: Machine Learning in Python’, Journal of Machine Learning Research, 12(85), pp. 2825–2830.

Peng, L. et al. (2015) ‘Large-scale RNA-Seq Transcriptome Analysis of 4043 Cancers and 548 Normal Tissue Controls across 12 TCGA Cancer Types’, Scientific Reports, 5(1), p. 13413. doi:10.1038/srep13413.

Putten, J.P.M. van and Strijbis, K. (2017) ‘Transmembrane Mucins: Signaling Receptors at the Intersection of Inflammation and Cancer’, Journal of Innate Immunity, 9(3), pp. 281–299. doi:10.1159/000453594.

R Core Team (2020) R: A language and environment for statistical computing. Vienna, Austria: R Foundation for Statistical Computing. Available at: https://www.r-project.org/ (Accessed: 16 February 2021).

Raffone, A. et al. (2019) ‘TCGA molecular groups of endometrial cancer: Pooled data about prognosis’, Gynecologic Oncology, 155(2), pp. 374–383. doi:10.1016/j.ygyno.2019.08.019.

Ritter, A. et al. (2018) ‘Diagnostic potential of micro RNAs expression profiles in serum and urine of breast and gynecologic cancer patients’, in. 38. Jahrestagung der Deutschen Gesellschaft für Senologie, Stuttgart, p. s-0038-1651787. doi:10.1055/s-0038-1651787.

Routledge, D. and Scholpp, S. (2019) ‘Mechanisms of intercellular Wnt transport’, Development, 146(10), p. dev176073. doi:10.1242/dev.176073.

Saraiya, M. et al. (2015) ‘US Assessment of HPV Types in Cancers: Implications for Current and 9-Valent HPV Vaccines’, JNCI Journal of the National Cancer Institute, 107(6), p. djv086. doi:10.1093/jnci/djv086.

Saso, S. et al. (2011) ‘Endometrial cancer’, BMJ, 343, p. d3954. doi:10.1136/bmj.d3954.

Schölkopf, B., Platt, J. and Hofmann, T. (eds) (2007) ‘Sparse Multinomial Logistic Regression via Bayesian L1 Regularisation’, in *Advances in Neural Information Processing Systems 19*. The MIT Press. doi:10.7551/mitpress/7503.003.0031.

Schüler-Toprak, S., Seitz, S. and Ortmann, O. (2017) ‘Hormones and risk of breast and gynecological cancer: A systematic review’, Der Gynäkologe, 50(1), pp. 43–54. doi:10.1007/s00129-016-4004-0.

Siegel, R.L. et al. (2021) ‘Cancer Statistics, 2021’, CA: A Cancer Journal for Clinicians, 71(1), pp. 7–33. doi:10.3322/caac.21654.

Simon, N. et al. (2011) ‘Regularization Paths for Cox’s Proportional Hazards Model via Coordinate Descent’, Journal of Statistical Software, 39(5). doi:10.18637/jss.v039.i05.

South, K. and Lane, D.A. (2018) ‘ADAMTS-13 and von Willebrand factor: a dynamic duo’, Journal of thrombosis and haemostasis: JTH, 16(1), pp. 6–18. doi:10.1111/jth.13898.

Steck, H. et al. (2007) ‘On Ranking in Survival Analysis: Bounds on the Concordance Index’, in Proceedings of the 20th International Conference on Neural Information Processing Systems. NIPS, pp. 1209–1216.

Tomczak, K., Czerwińska, P. and Wiznerowicz, M. (2015) ‘The Cancer Genome Atlas (TCGA): an immeasurable source of knowledge’, Contemporary Oncology, 19(1A), pp. A68–A77. doi:10.5114/wo.2014.47136.

Torre, L.A. et al. (2018) ‘Ovarian Cancer Statistics, 2018’, CA: a cancer journal for clinicians, 68(4), pp. 284–296. doi:10.3322/caac.21456.

Travaglino, A. et al. (2020) ‘Impact of endometrial carcinoma histotype on the prognostic value of the TCGA molecular subgroups’, Archives of Gynecology and Obstetrics, 301(6), pp. 1355– 1363. doi:10.1007/s00404-020-05542-1.

Valdez, J. et al. (2017) ‘On-demand dissolution of modular, synthetic extracellular matrix reveals local epithelial-stromal communication networks’, Biomaterials, 130, pp. 90–103. doi:10.1016/j.biomaterials.2017.03.030.

VanderVorst, K. et al. (2018) ‘Cellular and molecular mechanisms underlying planar cell polarity pathway contributions to cancer malignancy’, Seminars in Cell & Developmental Biology, 81, pp. 78–87. doi:10.1016/j.semcdb.2017.09.026.

Veena, M.S. et al. (2008) ‘Inactivation of the cystatin E/M tumor suppressor gene in cervical cancer’, Genes, Chromosomes & Cancer, 47(9), pp. 740–754. doi:10.1002/gcc.20576.

Vos, M.C. et al. (2016) ‘Limited independent prognostic value of MMP-14 and MMP-2 expression in ovarian cancer’, Diagnostic Pathology, 11(1), p. 34. doi:10.1186/s13000-016-0485-3.

Wang, M. et al. (2018) ‘A gene interaction network-based method to measure the common and heterogeneous mechanisms of gynecological cancer’, Molecular Medicine Reports [Preprint]. doi:10.3892/mmr.2018.8961.

Wang, Q. et al. (2018) ‘Unifying cancer and normal RNA sequencing data from different sources’, Scientific Data, 5(1), p. 180061. doi:10.1038/sdata.2018.61.

Wardwell-Swanson, J. et al. (2020) ‘A Framework for Optimizing High-Content Imaging of 3D Models for Drug Discovery’, SLAS DISCOVERY: Advancing the Science of Drug Discovery, 25(7), pp. 709–722. doi:10.1177/2472555220929291.

Whitley, B.R. et al. (2004) ‘Expression of active plasminogen activator inhibitor-1 reduces cell migration and invasion in breast and gynecological cancer cells’, Experimental Cell Research, 296(2), pp. 151–162. doi:10.1016/j.yexcr.2004.02.022.

Williams, E. et al. (2017) ‘Loss of polarity alters proliferation and differentiation in low-grade endometrial cancers by disrupting Notch signaling’, PLOS ONE, 12(12), p. e0189081. doi:10.1371/journal.pone.0189081.

Winkler, J. et al. (2020) ‘Concepts of extracellular matrix remodelling in tumour progression and metastasis’, Nature Communications, 11(1), p. 5120. doi:10.1038/s41467-020-18794-x.

Wu, B., Crampton, S.P. and Hughes, C.C.W. (2007) ‘Wnt signaling induces matrix metalloproteinase expression and regulates T cell transmigration’, Immunity, 26(2), pp. 227– 239. doi:10.1016/j.immuni.2006.12.007.

Wu, T. et al. (2021) ‘clusterProfiler 4.0: A universal enrichment tool for interpreting omics data’, The Innovation, 2(3). doi:10.1016/j.xinn.2021.100141.

Yadav, V.K. et al. (2020) ‘Computational analysis for identification of the extracellular matrix molecules involved in endometrial cancer progression’, PLoS ONE, 15(4). doi:10.1371/journal.pone.0231594.

Yang, M. et al. (2018) ‘Wnt signaling in cervical cancer?’, Journal of Cancer, 9(7), pp. 1277– 1286. doi:10.7150/jca.22005.

Yip, A.M. and Horvath, S. (2007) ‘Gene network interconnectedness and the generalized topological overlap measure’, BMC Bioinformatics, 8(1), p. 22. doi:10.1186/1471-2105-8-22.

Zhan, T., Rindtorff, N. and Boutros, M. (2017) ‘Wnt signaling in cancer’, Oncogene, 36(11), pp. 1461–1473. doi:10.1038/onc.2016.304.

Zhang, B. and Horvath, S. (2005) ‘A general framework for weighted gene co-expression network analysis’, Statistical Applications in Genetics and Molecular Biology, 4, p. Article17. doi:10.2202/1544-6115.1128.

Zhang, C. et al. (2020) ‘3D culture technologies of cancer stem cells: promising ex vivo tumor models’, Journal of Tissue Engineering, 11. doi:10.1177/2041731420933407.

Zhang, X. and Wang, Y. (2019) ‘Identification of hub genes and key pathways associated with the progression of gynecological cancer’, Oncology Letters [Preprint]. doi:10.3892/ol.2019.11004.

Zhao, J. et al. (2019) ‘The role of interleukin-17 in tumor development and progression’, Journal of Experimental Medicine, 217(1). doi:10.1084/jem.20190297.

Zou, H. and Hastie, T. (2005) ‘Regularization and variable selection via the elastic net’, Journal of the Royal Statistical Society: Series B (Statistical Methodology), 67(2), pp. 301–320. doi:10.1111/j.1467-9868.2005.00503.x.

