## Supplemental Figures and Tables for "Characterizing the extracellular matrix transcriptome of cervical, endometrial, and uterine cancers"

### **Supplementary Figures**

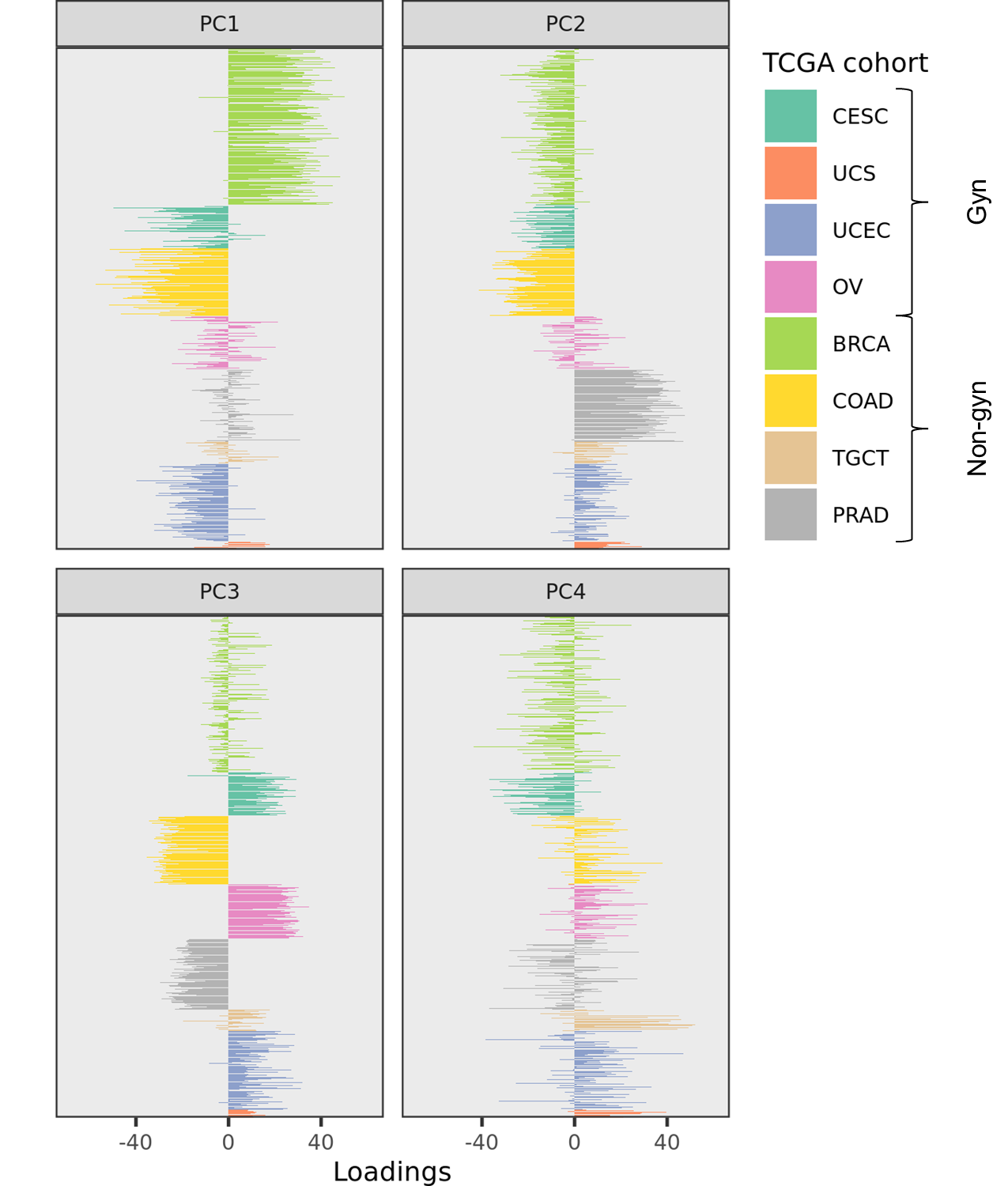

#### **Supplementary Figure 1. First 4 principal components of combined gynecological and non-gynecological TCGA cancer gene expression data.**

Loadings for first 4 principal components of the gynecological (gyn) and non-gynecological (non-gyn) cohorts investigated in our study.

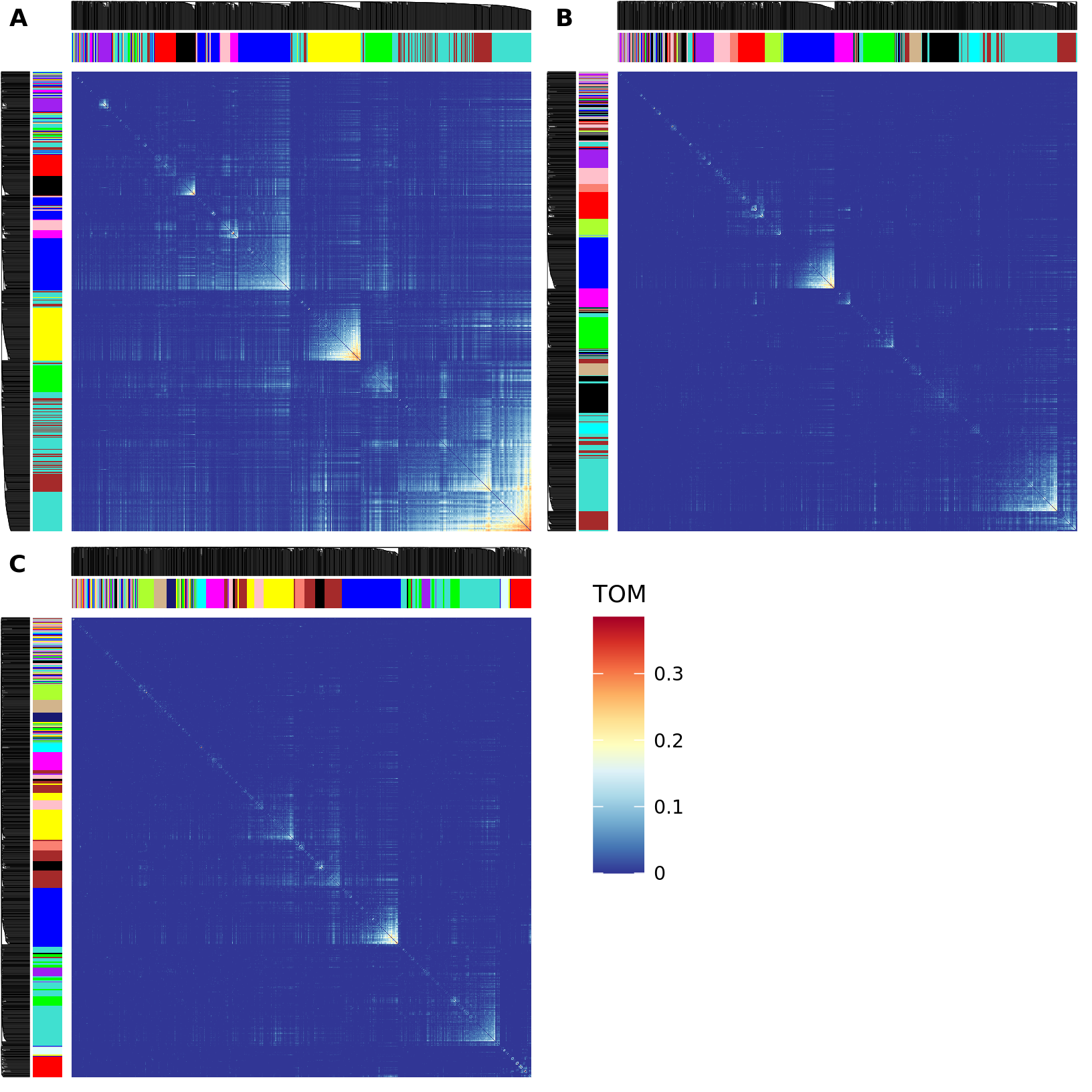

#### **Supplementary Figure 2. ECM topological overlap measure comparison.**

Visualization of topological overlap measure (TOM) matrices for **(A)** CESC, **(B)** UCEC, and **(C)** UCS cohorts. Dendrograms (black lines on $x$ and $y$ axes) and WGCNA module assignments (colors on $x$ and $y$ axes). Heatmap color represents gene-wise TOM (falling in the range $\left[ 0,1 \right]$). Gene-wise TOM estimates gene co-expression, so blue cells represent no co-expression while red cells represent stronger co-expression. Large pockets of yellow, orange, and red represent a large module, while smaller pockets represent a smaller module.

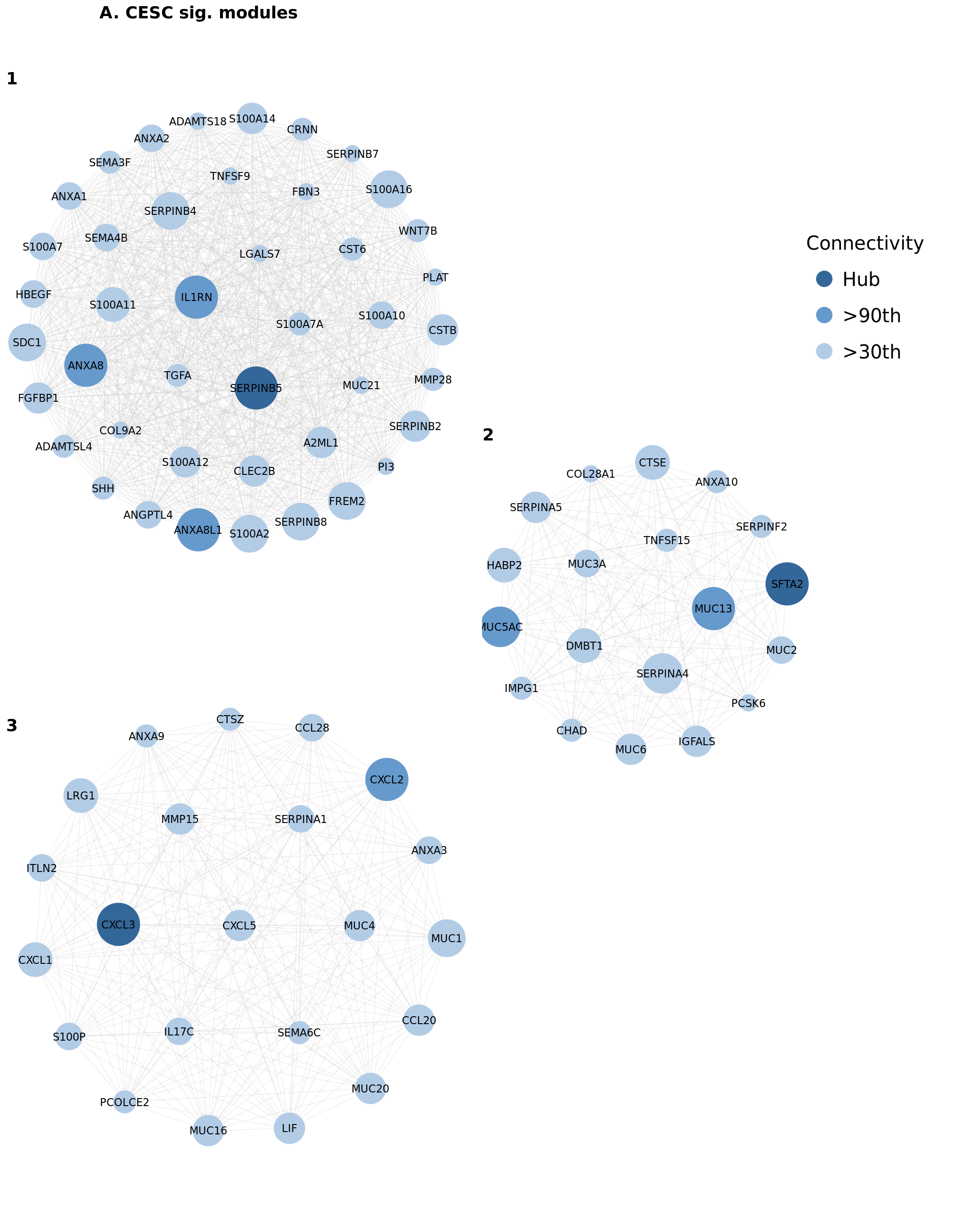

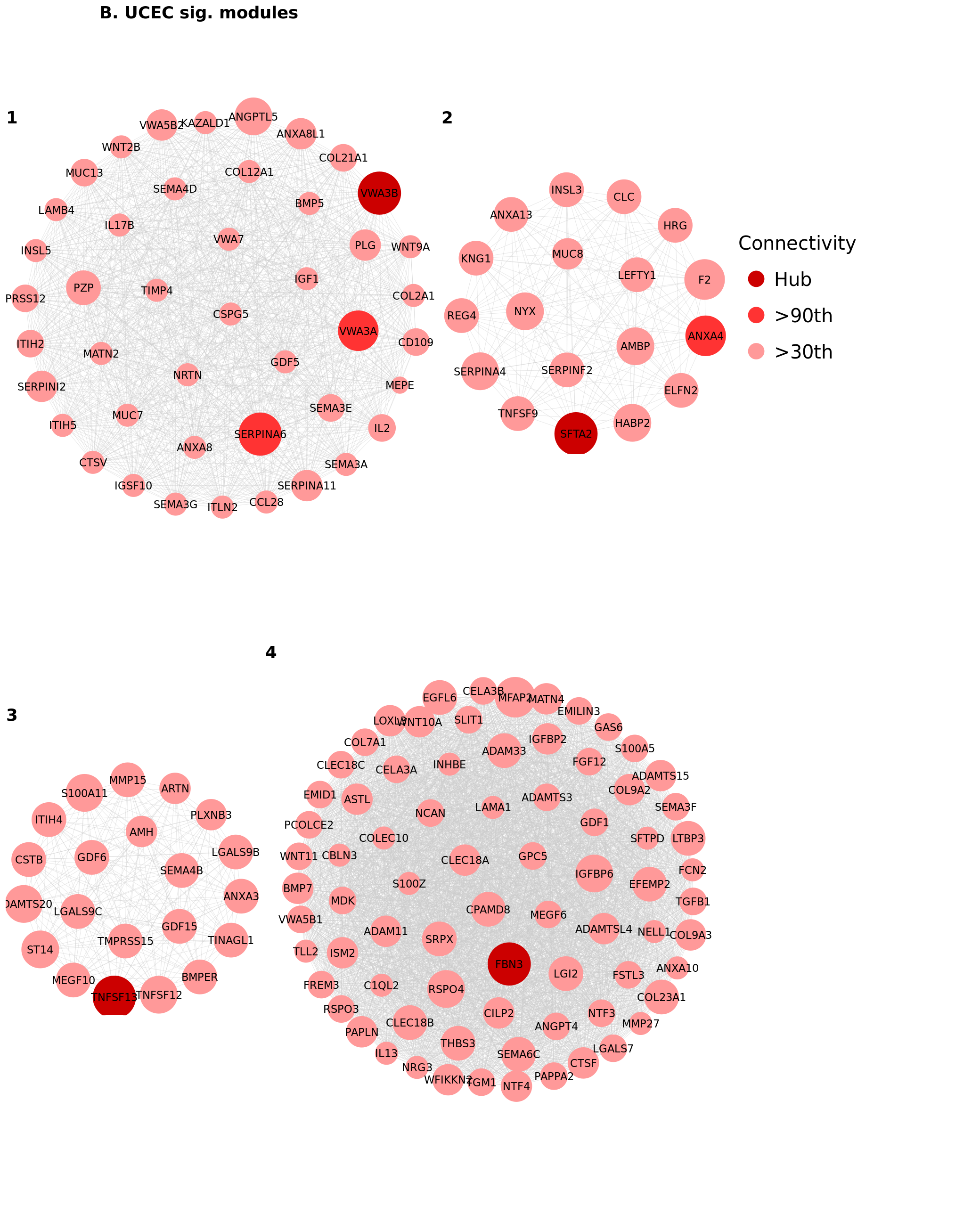

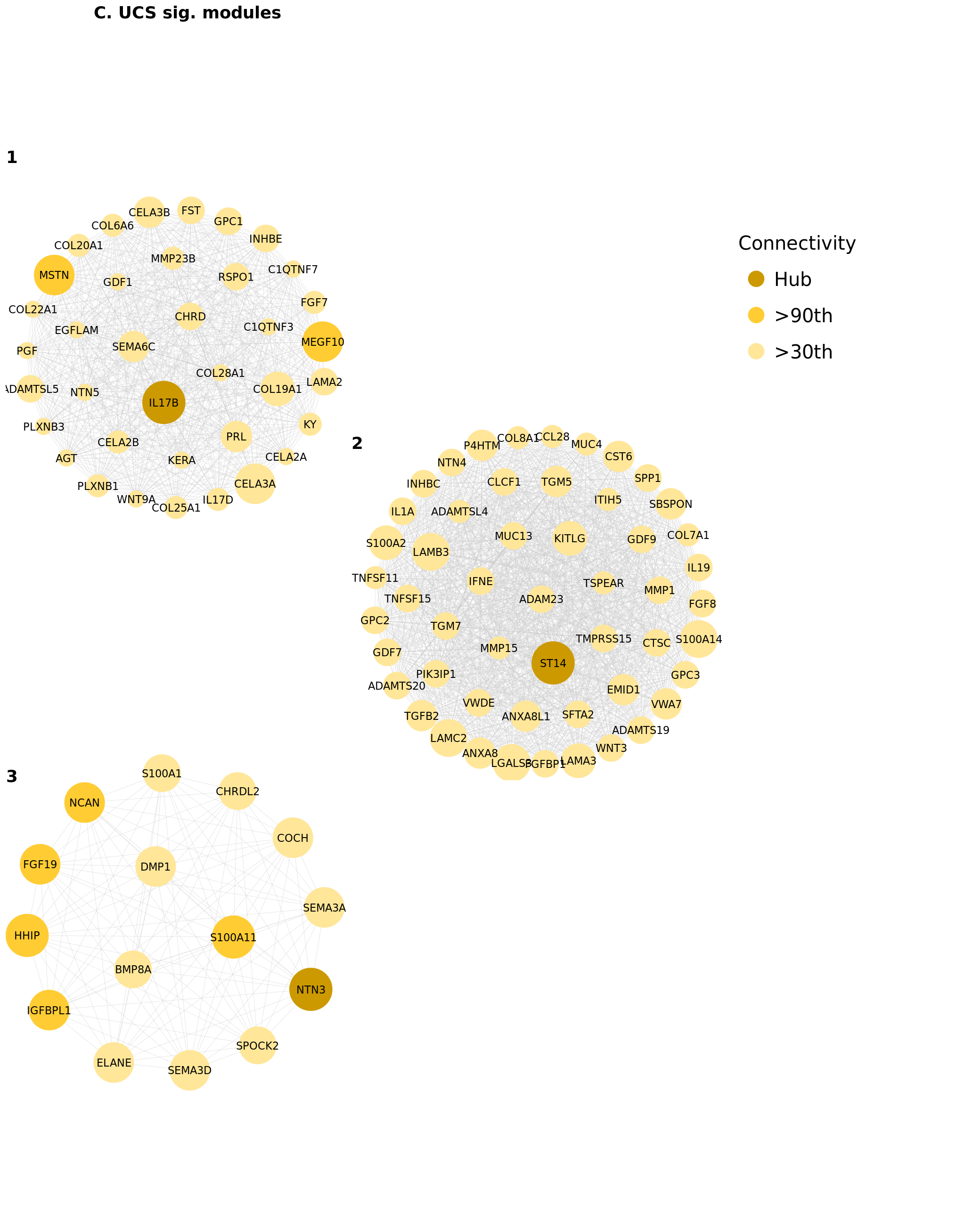

#### **Supplementary Figure 3. Filtered versions of all significant WGCNA modules for each cancer cohort.**

Full listing of significant WGCNA-assigned modules which were significantly correlated with one or more FIGO stage in **(A)** CESC, **(B)** UCEC, and **(C)** UCS. Module genes consist of differentially expressed matrisome genes (DEMGs) which were significant within their given module. These genes were filtered, scaled, and shaded based on their connectivity as compared to the connectivity of their module’s hub gene, the most connected DEMG in the module. Module DEMGs which were below the 30^th^ percentile of connectivity are not pictured, though they were utilized in our analyses. Module DEMGs which were in the 90^th^ percentile of connectivity are shaded darker than those below the 90^th^ percentile. Hub genes are shaded darkest. Connectivity was determined by the row-wise (gene-wise) sum of a given module’s adjacency matrix. Connectivity is relative to each module within each cohort, so node sizes cannot be compared between modules within the same or different cancer cohorts.

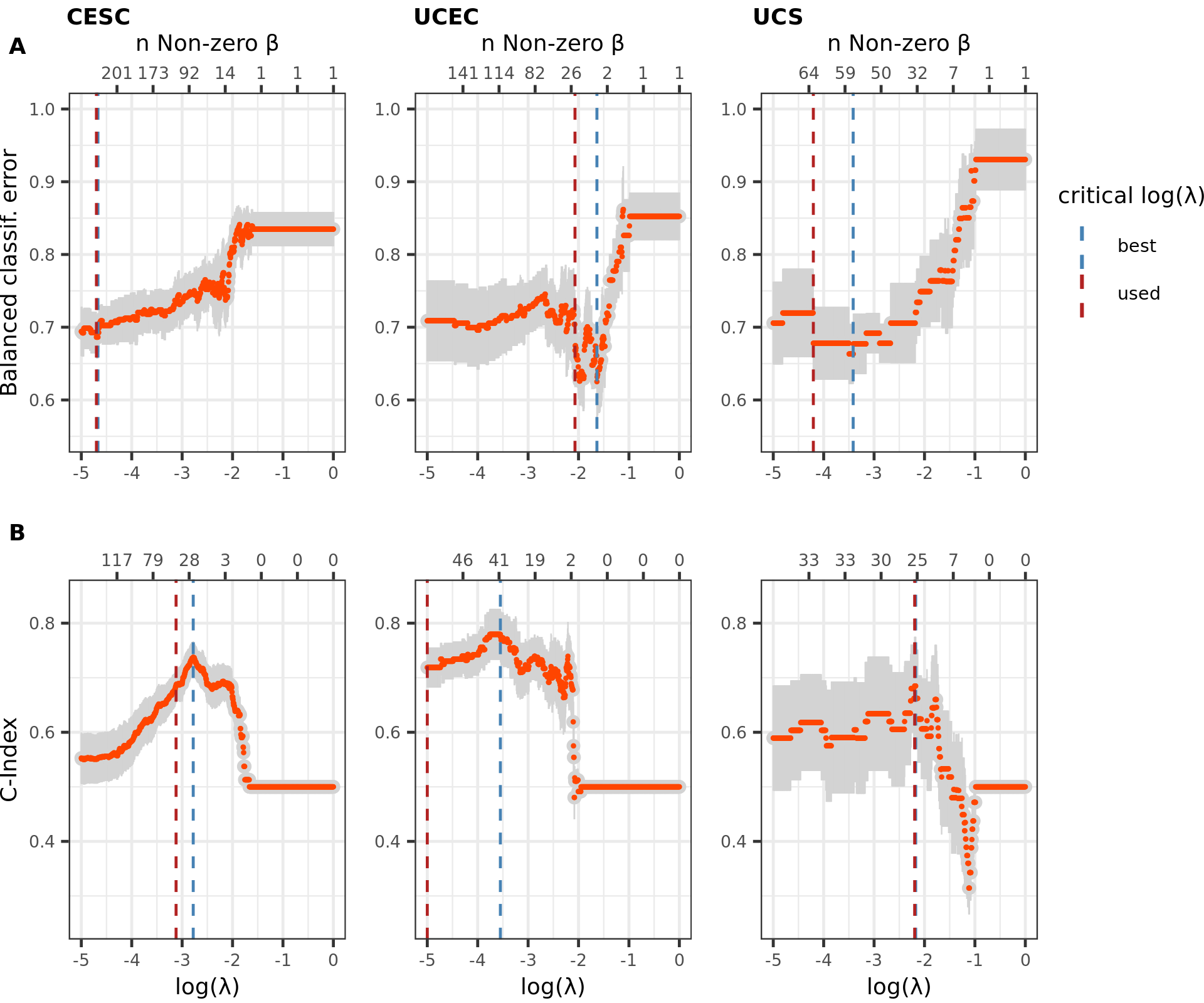

#### **Supplementary Figure 4. Penalized multivariable model selection for FIGO and survival models.**

Regularization strength ($\lambda$) tuning using grid search, applied to **(A)** multinomial regression for FIGO stage prediction and **(B)** Cox PH modeling for survival prediction. The performance of the worst-best multinomial regression model (CESC) and Cox PH model (UCS) were used to perform model selection in the other cohorts. Number of non-zero coefficients, $\beta$ (shown on top $x$-axis), tends to decrease as $\lambda$ increases so the first model (moving left-to-right) to achieve a target performance is often the least parsimonious model which can do so. The blue line represents the most performant model. The dotted red line represents the first model which achieved the benchmark performance for a given metric.

### **Supplementary Tables**

|  | ***Matrisome category*** (% DE) | | | | | |
| --- | --- | --- | --- | --- | --- | --- |
| **Cohort** | **Collagens** | **ECM glycoproteins** | **ECM regulators** | **ECM-affiliated proteins** | **Proteoglycans** | **Secreted factors** |
| CESC | 70% | 61% | 56% | 57% | 71% | 58% |
| UCEC | 67% | 63% | 61% | 59% | 69% | 60% |
| UCS | 74% | 64% | 56% | 50% | 74% | 59% |

#### **Table S1. Differential gene expression percentages by matrisome category.**

For each gynecological cancer cohort, percent of genes which were differentially expressed (DE) in each matrisome category.

| **Cohort** | $\boldsymbol{n}$ | $\boldsymbol{n}_{\boldsymbol{tumor}}$ | $\boldsymbol{n}_{\boldsymbol{normal}}$ | **Balanced accuracy** |
| --- | --- | --- | --- | --- |
| CESC | 272 | 259 | 13 | 1.000 |
| UCEC | 246 | 141 | 105 | 0.988 |
| UCS | 152 | 47 | 105 | 1.000 |

#### **Table S2. Predictive power of ECM cancer classification models for each gynecological cancer type.**

Performance of tumor/normal tissue stratification elastic net logistic regression models in each gynecological cancer cohort. A balanced accuracy of 1 represents a model which correctly classified all observations.

|  | ***Stage model sig. DEMGs*** | | | ***Survival model sig. DEMGs*** | | |
| --- | --- | --- | --- | --- | --- | --- |
| **Cohort** | **Point-biserial correlation** | **FIGO stage pairwise DGE analysis** | **L^1^ penalized multinomial regression** | **Censored time screening** | **Univariable KM/Cox PH** | **L^1^ penalized Cox PH regression** |
| CESC | 1 | 66 | 105 | 20 | 19 | 20 |
| UCEC | 3 | 61 | 13 | 0 | 1 | 26 |
| UCS | 0 | 33 | 38 | 0 | 2 | 16 |
|  | ***Set union of methods (total unique sig. DEMGs)*** | | | | | |
| CESC | 153 | | | 33 | | |
| UCEC | 70 | | | 26 | | |
| UCS | 62 | | | 18 | | |

#### **Table S3. Summary of model significant DEMG findings within each gynecological cancer cohort.**

Results of uni/multivariable analysis of matrisome genes with respect to FIGO stage and patient survival. FIGO stage significance metrics include 1) point-biserial correlation significance determined by Student $q$-value ($q<0.05$), 2) FIGO pairwise differential expression (DE) determined by Benjamini-Hochberg adjusted $p$-value and log fold-change ($p_{\text{adj}}<0.01$ and $abs\left( {log}_{2} \text{fold change} \right)>1$), and 3) L^1^ penalized multinomial regression significance determined by coefficient magnitude ($\beta_{\mathrm{gene}}\neq0$). Survival significance metrics include 1) censored time screening determined by log-rank test ($q<0.05$), Univariable KM/Cox PH significance determined by ($q<0.05$), and L^1^ penalized multivariable Cox PH regression significance determined by coefficient magnitude ($\beta_{\mathrm{gene}}\neq0$).

|  | ***Quantile*** | | | | |
| --- | --- | --- | --- | --- | --- |
| **Cohort** | **0%** | **25%** | **50%** | **75%** | **100%** |
| CESC | 0.002 | 0.008 | 0.013 | 0.02 | 0.072 |
| UCEC | 0 | 0.002 | 0.003 | 0.004 | 0.026 |
| UCS | 0 | 0.001 | 0.002 | 0.004 | 0.017 |

#### **Table S4. ECM Topological overlap measure comparison between gynecological cancers.**

Quantiles of mean gene-wise topological overlap measures (TOMs) for matrisome genes in each cancer cohort. CESC has far higher gene-wise average TOMs, implying more co-expression among matrisome genes in CESC than the other cancers.

| **Cohort** | **Significant modules** | **Stage network sig. DEMGs** |
| --- | --- | --- |
| CESC | 3 | 140 |
| UCEC | 4 | 159 |
| UCS | 3 | 129 |

#### **Table S5. Matrisome network analysis summary for all gynecological cancers.**

Number of stage significant modules and number of stage network significant DEMGs in each gynecological cancer cohort.

| **Cohort** | **DEMGs** | **Stage sig. DEMGs** | | | **Survival sig. DEMGs** | **Stage and survival DEMG overlap** |
| --- | --- | --- | --- | --- | --- | --- |
|  |  | **Model sig. DEMGs** | **Network sig. DEMGs** | **Model or network sig. DEMGs** | **Model sig. DEMGs** |  |
| CESC | 593 | 153 | 140 | 251 | 33 | 13 |
| UCEC | 618 | 70 | 159 | 207 | 26 | 9 |
| UCS | 595 | 62 | 129 | 177 | 18 | 3 |

#### **Table S6. Set meta-analysis for DEMGs and FIGO stage/survival significant DEMGs.**

Breakdown of significant genes by cohort and category. FIGO stage significant DEMGs are defined as DEMGs which are also stage model or network significant. Survival significant DEMGs are defined as DEMGs which are survival model significant. Full overlap includes DEMGs which are both stage significant and survival significant.

| **Matrisome Category** | **Master list count** | **Dataset count** |
| --- | --- | --- |
| Collagens | 44 | 43 |
| ECM Glycoproteins | 195 | 192 |
| ECM Regulators | 238 | 233 |
| ECM-affiliated Proteins | 171 | 168 |
| Proteoglycans | 35 | 35 |
| Secreted Factors | 344 | 337 |

#### **Table S7. Matrisome category counts according to full human matrisome master list.**

Gene counts within each matrisome category in the matrisome master list.

| **Cohort** | **Metrics** | | | |
| --- | --- | --- | --- | --- |
|  | **Best score** | $\boldsymbol{n}$ **DEMG coef. best model** | **Used score** | $\boldsymbol{n}$ **DEMG coef. used model** |
| ***FIGO stage model balanced classification error*** ($baseline=0.75$, smaller is better) | | | | |
| CESC | 0.686 | 102* | 0.686 | 105* |
| UCEC | 0.625 | 3 | 0.668 | 13 |
| UCS | 0.663 | 33 | 0.678 | 38 |
| ***Survival model concordance index*** ($baseline=0.5$, larger is better) | | | | |
| CESC | 0.738 | 10 | 0.687 | 20 |
| UCEC | 0.780 | 25 | 0.719 | 26 |
| UCS | 0.685 | 16* | 0.685 | 16* |

#### **Table S8. Penalized multivariable machine learning metrics for stage and prognosis prediction.**

Objective function scores and retained coefficients in each cohort for multivariable FIGO stage (L^1^ multinomial regressor) and survival (L^1^ Cox PH) models. The used model is simply the first model for which an acceptable performance (performance of the worst of the best models between the three cohorts) was achieved. FIGO baseline: guess majority class for all observations, survival baseline: random guess about patient survival times. The number of DEMG coefficients in each model is simply the number of retained coefficients (i.e., matrisome genes) after filtering based on DEMG status.

*CESC multinomial and UCS Cox PH performance were used to threshold the other models. Discrepancies between best/used for these models were due to three decimal place rounding used when thresholding model performance.
